# Trellis Single-Cell Screening Reveals Stromal Regulation of Patient-Derived Organoid Drug Responses

**DOI:** 10.1101/2022.10.19.512668

**Authors:** María Ramos Zapatero, Alexander Tong, Jahangir Sufi, Petra Vlckova, Ferran Cardoso Rodriguez, Callum Nattress, Xiao Qin, Daniel Hochhauser, Smita Krishnaswamy, Christopher J. Tape

**Author notes:** These authors contributed equally (María Ramos Zapatero, Alexander Tong).

## Abstract

Patient-derived organoids (PDOs) can model personalized therapy responses, however current screening technologies cannot reveal drug response mechanisms or study how tumor microenvironment cells alter therapeutic performance. To address this, we developed a highly-multiplexed mass cytometry platform to measure post translational modification (PTM) signaling in >2,500 colorectal cancer (CRC) PDOs and cancer-associated fibroblasts (CAFs) in response to clinical therapies at single-cell resolution. To compare patient- and microenvironment-specific drug responses in thousands of single-cell datasets, we developed *Trellis* — a highly-scalable, hierarchical tree-based treatment effect analysis method. Trellis single-cell screening revealed that on-target cell-cycle blockage and DNA-damage drug effects are common, even in chemorefractory PDOs. However, drug-induced apoptosis is patient-specific. We found drug-induced apoptosis does not correlate with genotype or clinical staging but does align with cell-intrinsic PTM signaling in PDOs. CAFs protect chemosensitive PDOs by shifting cancer cells into a slow-cycling cell-state and CAF chemoprotection can be reversed by inhibiting YAP.

**Highlights:**

- >2,500 single-cell PTM signaling, DNA-damage, cell-cycle, and apoptosis responses from drug-treated PDOs and CAFs.
- Trellis: hierarchical tree-based treatment effect method for single-cell screening analysis.
- PDOs have patient-specific drug responses that align with cell-intrinsic PTM signaling states.
- CAFs chemoprotect PDOs by altering PDO cell-state via YAP signaling.

## 1 Introduction

Tumors are heterogeneous cellular systems comprising cancer cells, stromal fibroblasts, and various immune cells. Tumor phenotypes are regulated by cell-intrinsic mutations within cancer cells and cell-extrinsic cues from the tumor microenvironment (TME) [1]. Colorectal cancer (CRC) kills >0.9 million people per year worldwide [2] and is characterized by a high inter-patient genetic heterogeneity and patient-specific responses to therapy [3]. Cancer associated fibroblasts (CAFs) are one of the most abundant cell-types in the CRC TME [4]. CAF abundance correlates with poor overall survival [5], and influences response to both targeted therapies [6] and radiotherapy [7]. However, there is a lack of understanding regarding how CAFs regulate cancer cell response to therapy and to what extent stromal regulation is patient-specific.

Patient-derived organoids (PDOs) are personalized cancer models [8] that can mimic their parent tumors’ response to chemotherapies [9] — with several studies proposing PDOs as personalized avatars of drug response [10]. However, epithelial PDO monocultures cannot model the influence of stromal cells on therapy response. PDOs can be co-cultured with stromal and immune cells to recapitulate elements of the TME [11], but how this alters PDO phenotypes and personalized drug response is unknown. Moreover, PDO drug sensitivity is typically measured using bulk live/dead viability assays [12] that cannot resolve cell-type-specific data from co-cultures and provide no mechanistic insight into drug responses [13].

To overcome these limitations, we developed a highly-multiplexed Thiol-reactive Organoid Barcoding *in situ* (TOBis) mass cytometry [14, 15] platform to study how anti-cancer therapies regulate the cell-state, DNA-damage response, and post-translational modification (PTM) signaling of CRC PDOs in the presence or absence of CAFs at single-cell resolution across >2,500 PDO-CAF cultures. To compare single-cell drug responses from thousands of cell-type-specific datasets, we developed *Trellis*, a hierarchical tree-based treatment effect analysis method that derives generalized optimal transport distances between samples after normalizing by their own controls. TOB*is* mass cytometry and Trellis revealed drug-induced PTM signaling responses are PDO-specific and demonstrated CAFs protect CRC cells from chemotherapy by shifting epithelial cells into a slow-cycling cell-state. CAF chemoprotection could be rationally reversed by inhibiting YAP-signaling using insights from single-cell PTM data, demonstrating the utility of PTM-focused drug screening for overcoming therapy resistance. These results illustrate the functional intertumoral heterogeneity of patient-specific drug response mechanisms and suggest TME cells should be included in future PDO models.

## 2 Results

### Patient- and Microenvironment-Specific Single-Cell PTM PDO-CAF Drug Analysis

To study how CAFs regulate patient-specific drug response signaling, we established a high-throughput 3D organoid co-culture system comprising 10 CRC PDOs [12] (Table S1) cultured either alone or with CRC CAFs [16, 17]. Organoid cultures were treated in triplicate with either vehicle control, or titrated combinations of clinical therapies fluoropyrimidine 5-fluorouracil (5-FU), SN-38 (active metabolite of Irinotecan), Oxaliplatin, and Cetuximab (EGFR inhibitor). The pre-clinical therapy LGK974 (PORCN inhibitor) [12] was also studied to investigate PDO-CAF WNT signaling and Berzosertib (VX-970) was included as ATR inhibition has been hypothesized to synergize with DNA-damaging agents in CRC [18] (Figure 1a) (Table S2). Following treatment, each culture was fixed, stained with thiol-reactive monoisotopic TOB*is* barcodes [15], pooled, dissociated into single-cells, stained with a panel of 44 rare-earth metal antibodies (spanning cell-type, cell-state, DNA-damage, and PTM signaling markers (Table S3)), and analyzed by mass cytometry (Figure 1b). Following multiplexed debarcoding [19] and cell-type-specific gating, we obtained >10 million PDO cells and >15 million CAFs from 2,520 3D cultures (3,360 cell-type-specific single-cell PTM signaling datasets).

**Figure 1:**
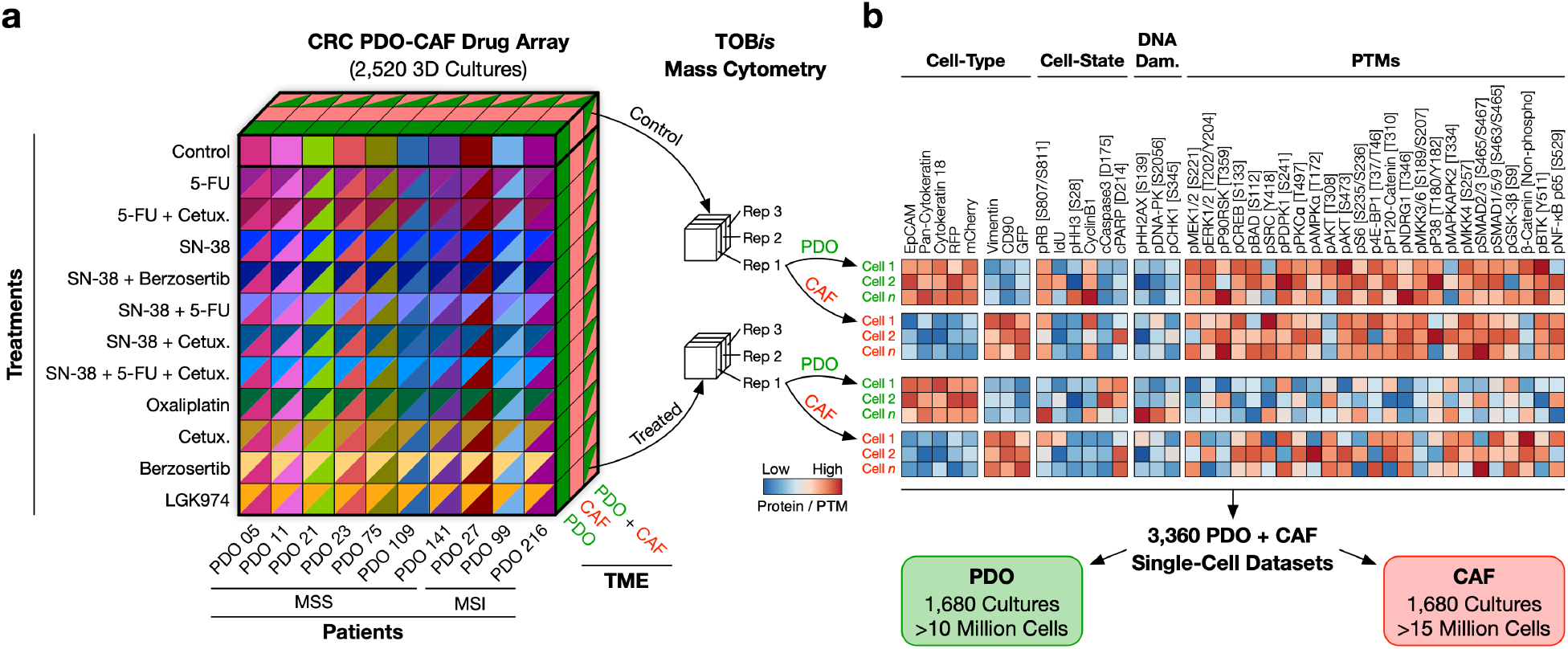
TOB*is* MC Single-Cell PTM PDO-CAF Drug Responses. **a)** Multidimensional array of 10 CRC PDO (7 microsatellite stable (MSS), 3 microsatellite instable (MSI)) treated with 11 titrated drug combinations either alone or in co-culture with CRC CAFs in triplicate (2,520 3D cultures). **b)** PDO-CAFs were barcoded *in situ* using TOBis, stained with 44 rare-earth metal antibodies spanning cell-type identification, cell-state, DNA-damage response, and PTM signaling, and analyzed by MC (3,360 single-cell PTM datasets).

### Trellis: Hierarchical Tree-Based Single-Cell Treatment Effect Analysis

Highly-multiplexed single-cell cytometry screening data presents several analytical challenges. First, existing work on large single-cell data uses the manifold structure of transcriptomic technologies, where cell distances are locally Euclidean [20–24]. However, in cytometry data antibody panels are designed based on prior biological knowledge, and analyzed using gating strategies that follow a hierarchical structure, which are better described by tree distances rather than a single smooth manifold. Second, our PDO-CAF PTM screening data contains >2,500 conditions with >25 million cells. Existing state of the art to analyze such large datasets is to compare cluster proportions between single-cell samples [25–27]. Emerging methods can compare distributions using earth mover’s distance (EMD), but only at course granularity [22], or by using graph diffusion which does not account for the hierarchical tree structure of cytometry data [23]. As highly-multiplexed single-cell screening datasets are becoming increasingly common, there is a need for tools that can efficiently compare thousands of single-cell conditions. Finally, large screening datasets compare independent systems (e.g. patients, microenvironments, and/or technical batches) perturbed by constant treatments. For this, internal controls need to be leveraged, such that multiple controls and treatments can be directly compared in a common computational space. To solve these problems, we developed *Trellis*.

Comparisons between single-cell datasets typically treat all markers equally — irrespective of prior biological knowledge. While equal weighting may be appropriate for unbiased single-cell methods such as scRNA-Seq, Trellis leverages the experimental design of cytometry data using a ‘branch’ tree hierarchy of well-defined biological processes (e.g. cell-type hierarchies or cell-states) that supervenes upon randomized ‘leaves’ of latent biological significance (via four levels of four k-means clusters) (Figure 2a). This enables an automated assessment of cell populations that mimics human intuition in the design of the experiment, and subsequently its interpretation (Figure S1). Trellis can leverage any gating strategy that returns a single hierarchy or multiple hierarchies (Algorithm 1 line 3).

#### Algorithm 1: High-level Trellis algorithm for comparing single-cell treatment effects

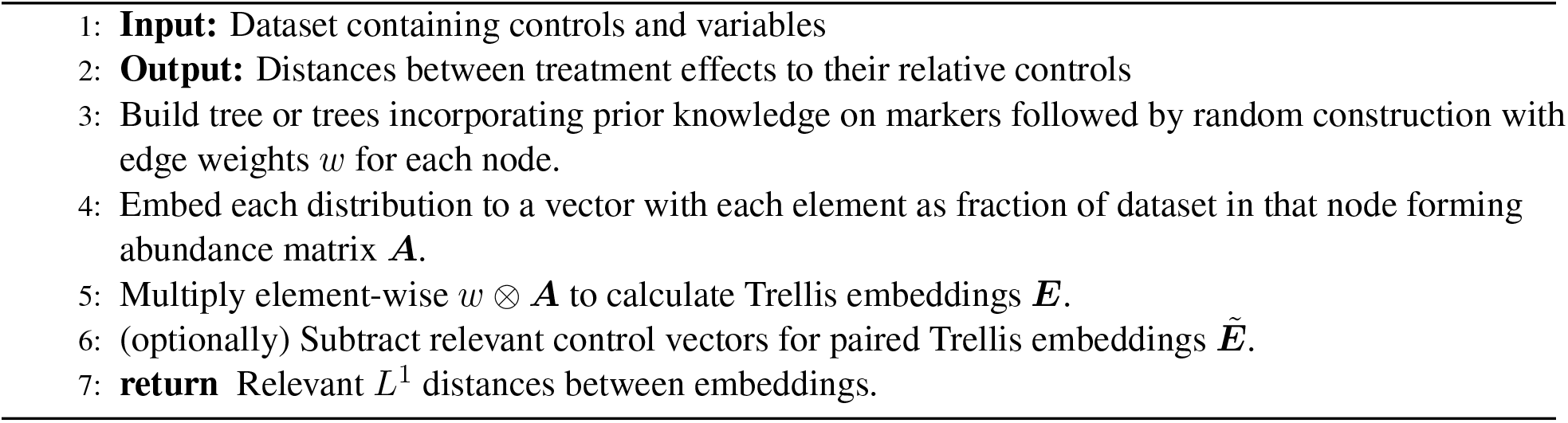

**Figure 2:**
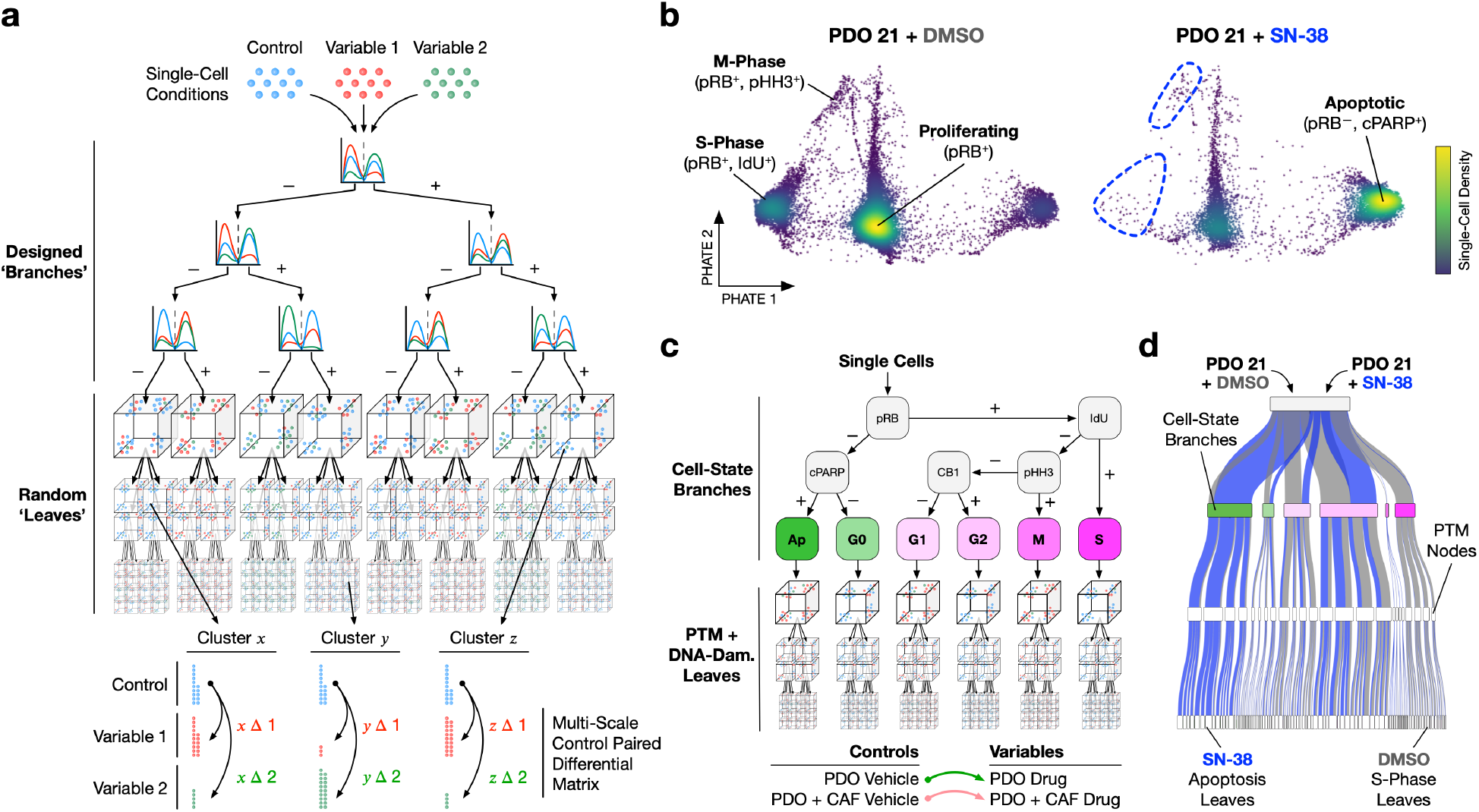
Trellis Single-Cell Treatment Effect Analysis. **a)** Single cells from control and variable conditions are distributed through a tree comprising designed ‘branches’ that supervene upon randomized *k*-means clustering nodes, ending in ‘leaves’. Branches weigh hierarchical gating strategies while nodes and leaves leverage latent parameters. In each node of the tree, variables are subtracted from paired controls to create a multi-scaled differential matrix (representing a Kantorovich-Rubinstein norm) that scales to thousands of conditions. **b)** Single-cell density PHATEs of PDO 21 treated with DMSO or SN-38 (irinotecan). SN-38 results in cell-cycle exit (IdU^−^, pHH3^−^, and pRB^−^) and induction of apoptosis (cPARP^+^). **c)** Trellis hierarchy for single-cell PDO on-target drug responses leveraging cell-state branches and randomized PTM and DNA-damage parameters. Trellis scores are calculated per PDO by comparing untreated controls to drugs for both mono-cultures and co-cultures. CB1, Cyclin B1. **d)** Sankey diagram showing data from b) distributing through the Trellis layout in c) (terminal leaves not shown).

Once one or multiple hierarchies are defined, Trellis then embeds each single-cell distribution into a vector such that for two distributions, the L^1^ distance between embeddings is equivalent to the EMD between the two distributions along the defined tree or forest (Algorithm 1 lines 4-5). By reducing a complicated and inefficient distance calculation to a vector distance, Trellis can scale to larger datasets by leveraging existing work in high-dimensional distance computation (Figure S2). For instance, if we only need to find the nearest neighbor treatments for non-linear embedding [21, 28], we leverage fast nearest neighbor algorithms such as KD-trees as used in PHATE [21], Annoy [29] used in UMAP [30] and Scanpy [31], or locally sensitive hashing [32, 33].

As single-cell screens increase in size and complexity, the use of internal controls enables the comparison of independent variables in parallel. Existing distribution comparison methods cannot easily incorporate pairing of controls to variables, indeed EMD is not even defined for the difference of distributions. To solve this, Trellis can easily be extended to ‘paired’ Trellis (Algorithm 1 line 6), where paired controls are subtracted from treatment samples to directly compare treatment effects. We prove this is equivalent to a Kantorovich-Rubenstein (KR) norm with tree ground distance (Prop. 2). This KR norm cannot be computed with standard Wasserstein distance methods (even for small problems [34, 35]) but can be calculated by Trellis. Paired Trellis therefore enables thousands of variables to be compared to their internal controls in a common computational space — enabling clear distinction of individual treatment effects in paralleled high-dimensional single-cell screening data (Figure S1a).

In summary, Trellis uses a prior-driven tree domain to compute the generalized Wasserstein distance between thousands of single-cell samples. Pairing treatments to controls enables paralleled visualization of treatment effects (Figure S1a) and reduces batch effects in serially acquired screening data (Figure S1b). Prior-driven branches further resolve biologically important treatment effects (Figure S1c). Trellis outperforms existing single-cell treatment effect methods (Figure S2a) and the tree domain structure enables thousands of single-cell datasets to be analyzed rapidly (Figure S2b). Prior-driven branches are customizable to different biological questions and Trellis recapitulates features of published datasets (Figure S3). Further detail on Trellis’ scalability, theoretical soundness, and robustness can be found in Methods.

### Trellis Single-Cell Analysis of PDO Cell-State and PTM Signaling

Anti-cancer drugs typically induce major shifts in cell-cycle and apoptosis that can be detected by mass cytometry. For example, SN-38 inhibits topoisomerase 1 [36], resulting in S-phase blockage, cell-cycle exit, and induction of apoptosis (Figure 2b). Similarly, 5-FU blocks nucleotide biosynthesis by inhibiting thymidylate synthase [37] which subsequently stalls S-phase entry, whereas oxaliplatin induces ribosome biogenesis stress to block mitotic progression [38]. Capturing shifts in cell-state is therefore crucial for understanding on-target drug responses in single-cell data.

In mass cytometry, cell-state is identified by hierarchical gating of pRB, IdU, pHH3, Cyclin B1, and cPARP/cCaspase3 [39, 40] and is therefore well suited for Trellis branches. For cell-type-specific analysis of PDO-CAF co-cultures we designed a Trellis hierarchy using cell-state-driven branches that supervene upon randomized DNA-damage and PTM signaling leaves (Figure 2c) (Figure S4a). This tree topology sensitizes Trellis to canonical on-target drug-induced shifts in cell-cycle and apoptosis while also leveraging latent changes in DNA-damage and PTM signaling (Figure 2d) (Figure S4b-e).

### Trellis Analysis of Cell-Type-Specific PDO-CAF Drug Responses

We used Trellis to analyze 3,360 (1,680 PDO, 1,680 CAF) single-cell PTM profiles (>25 million single-cells) (Figure 3a) in order to explore drug-, patient-, and microenvironment-specific therapy responses for both PDOs (Figure 3b-d) and CAFs (Figure S5). Since Trellis performs pairwise normalization to internal controls, all controls group on the left side of the graph (Figure 3b) (Figure S1a) and each treatment embeds relative to their controls, depending on their distribution through the Trellis tree. This enables therapeutic effects to be visualized across PHATE 1 and mechanistic response in PHATE 2 (Figure S6). If the same drug were to have have an equal effect on all PDOs, Trellis would group each condition by drug type. However, Trellis revealed PDO treatment effects are characterized not by drug type, but by patient-specific signaling responses.

**Figure 3:**
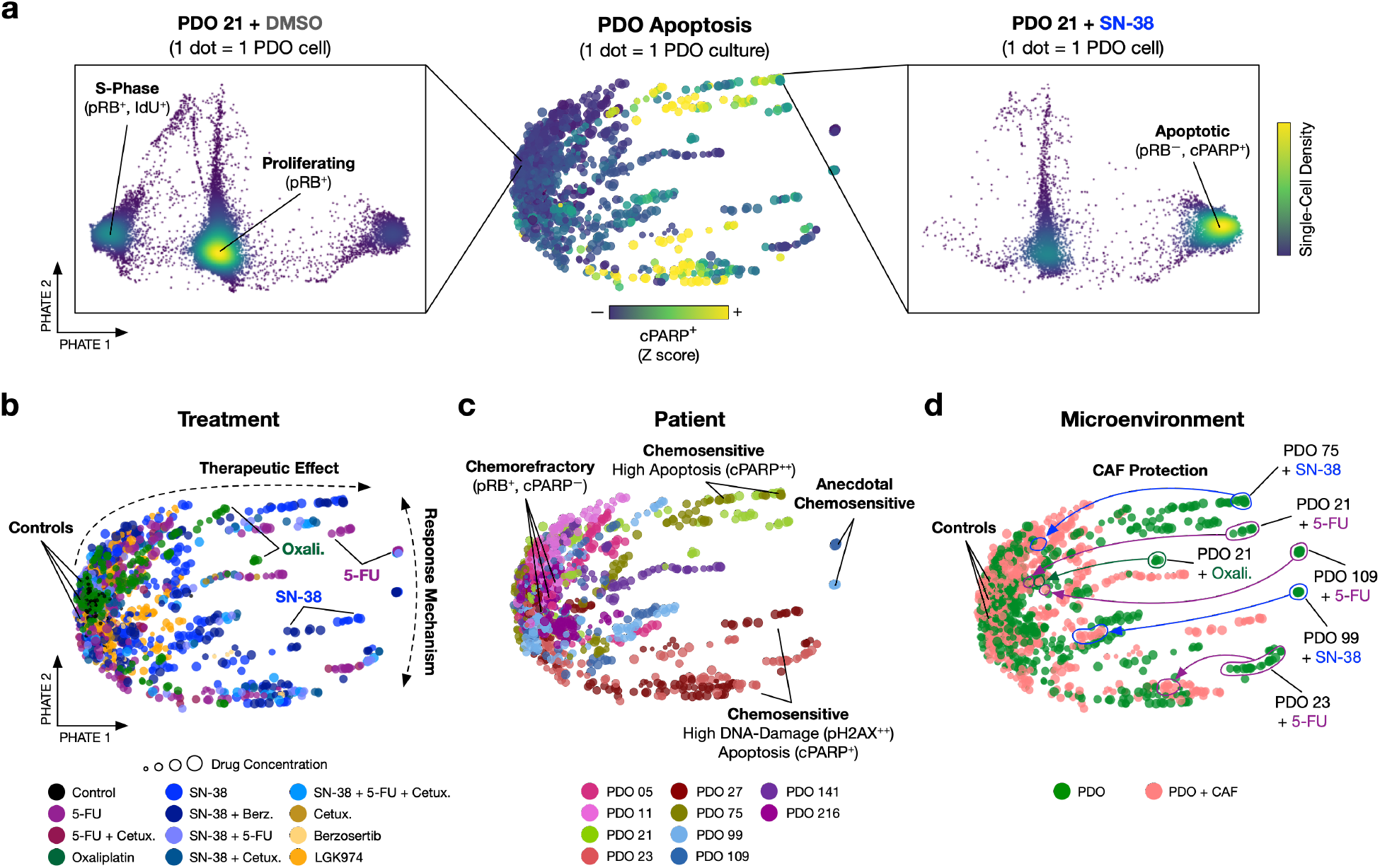
Trellis Analysis of Single-Cell PDO-CAF Drug Responses. **a)** Trellis-PHATE of 1,680 PDO single-cell PTM profiles (1 dot = 1 organoid culture comprising >5,000 single-cells) colored by apoptosis with representative single-cell density embeddings of PDO 21 + DMSO or + SN-38. **b)** PDO drug treatment-specific responses. Controls group on the left, with treatment effects spreading across PHATE 1 and response mechanisms resolving across PHATE 2. **c)** Patient-specific drug responses illustrate different chemosensitive mechanisms and chemorefractory patients. **d)** CAFs provide patient-specific chemoprotection from 5-FU, SN-38, and oxaliplatin.

We observed four patient-grouped responses to 5-FU, SN-38, and oxaliplatin chemotherapies: 1) broadly chemosensitive with high apoptosis (PDOs 21 and 75), 2) broadly chemosensitive with apoptosis and a strong DNA-damage response (PDOs 23 and 27), 3) anecdotally chemosensitive (i.e. only apoptotic with a specific drug) (PDOs 99 and 109), and 4) chemorefractory with minimal apoptosis and low DNA-damage response (PDOs 05, 11, 141, and 216). Cetuximab, Berzosertib, and LGK974 generally had modest effects on PDO cell-state and PTMs relative to chemotherapies (Figure 3b) (Figure S6). While PDOs demonstrate clear patient- and microenvironment-specific drug responses, CAF signaling does not cluster by patient or drug (Figure S5), suggesting chemotherapies mainly alter the cell-state, DNA-damage, and PTM profiles of PDOs, not CAFs. Trellis further revealed CAFs protect some PDOs from chemotherapies (Figure 3d).

### PDO Drug Signaling Responses Are Patient-Specific

PDOs have been proposed as personalized avatars of drug response [10], but how clinical treatments mechanistically alter patient-specific PDO biology is not well understood. To explore patient-specific drug response signaling, we updated the designed branches of the Trellis tree by combining cell-state parameters with a pHH2AX [S139] detection layer to enrich on-target DNA double-strand breaks and analyzed each patient drug response in parallel (Figure S7a-d). Patients continue to display either broad (PDOs 21, 23, 27, and 75) or anecdotal (PDOs 99 and 109) chemotherapeutic sensitivity, and multiple examples of drug insensitivity (Figure 4a).

**Figure 4:**
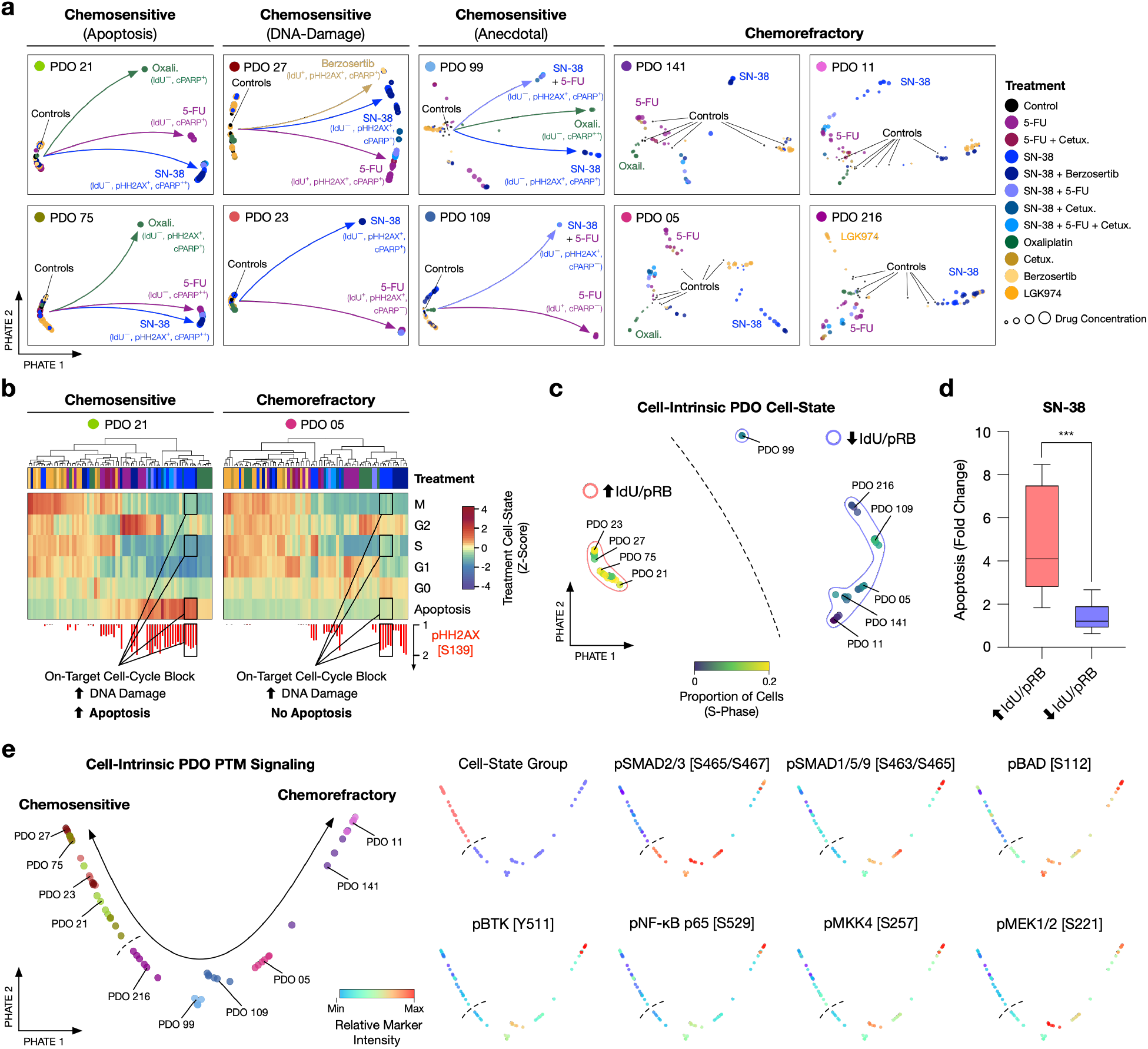
PDO Drug Response Mechanisms Are Patient-Specific and Align With Cell-Intrinsic Cell-State and PTM Signaling. **a)** Trellis-PHATE patient-specific PDO drug responses (840 single-cell PTM datasets). **b)** Patient-specific distribution of cells within Trellis branches reveals on-target cell-state shifts upon drug treatments. Treatment cell-state quantifies the fold change of the proportion of cells/cell state over the controls for each treatment (Z-score). DNA damage is quantified by the fold change of the proportion pHH2AX^+^ cells over the controls. **c)** Trellis-PHATE resolves high IdU/pRB (red outline) and low IdU/pRB (blue outline) cell-intrinsic cell-state PDO groups (colored by proportion of cells in S-phase). **d)** SN-38-induced apoptosis in low IdU/pRB and high IdU/pRB PDOs. Unpaired t-test, *** <0.001. **e)** TreEMD-PHATE of cell-intrinsic PTM signaling nodes demonstrates PTMs up-regulated in chemorefractory PDOs.

Unlike univariate live/dead metrics used in traditional drug screens, TOB*is* mass cytometry can detect on-target treatment effects that do not result in cell death. For example, SN-38 induces on-target S-phase blockage and double-strand breaks in both PDO 21 and PDO 05, yet only PDO 21 translates genotoxic stress into apoptosis (Figure 4b). Similarly, in PDOs 23 and 99, 5-FU and SN-38 result in a large DNA-damage response and stalled mitosis respectively, but no apoptosis (Figure S7e). 5-FU and SN-38 can clearly induce double-strand breaks and cell-cycle arrest in these PDOs, but they do not translate genotoxic replication stress into cell death. In fact, across nearly all PDOs tested, SN-38 (Figure S7f), oxaliplatin (Figure S7g), and 5-FU (Figure S7h) display on-target mitotic arrest (83%), but only a subset of patient and treatment combinations trigger apoptosis (40%). This suggests on-target drug responses are common in CRC PDOs, but often insufficient to induce cell death.

The patient-specific drug sensitivity demonstrated by several PDOs reinforces the notion that PDOs could be used to identify drugs uniquely potent to an individual’s cancer. For example, in PDO 99, 5-FU blocks mitosis and SN-38 causes a large DNA-damage response – yet neither chemotherapy induces substantial apoptosis. However, when treated with oxaliplatin, PDO 99 exits the cell-cycle and enters apoptosis (Figure S7e). Unlike 5-FU and SN-38, oxaliplatin does not kill cells directly through blocking S-phase, but via inducing ribosome biogenesis stress [38]. PDO 99 appears refractory to cytostatic stress but hypersensitive to ribosome biogenesis stress. Similarly, ATR inhibitors block single-stranded DNA-damage response and typically synergize with DNA-damage inducing drugs [18]. However, we find Berzosertib only increases SN-38-induced apoptosis in MSI PDOs (Figure S8), suggesting ATR inhibitors might only be effective in MSI patients.

### Chemosensitive PDOs Have Distinct Cell-Intrinsic PTM Signaling Profiles

We next sought to understand features common to chemosensitive and chemorefractory PDOs. Therapeutic response does not correlate with MSI/MSS status, clinical staging, anatomical location, *KRAS*, or *APC* genotypes (Figure S9a) (Table S1). However, baseline PDO cell-state and PTM signaling profiles are patient-specific and align with chemosensitivity (Figure 4c) (Figure S9b-d). Chemosensitive PDOs 21, 23, 27, and 75 are highly proliferative at baseline and experience canonical S-phase blockage, increased DNA-damage, and apoptosis when treated with both 5-FU and SN-38. In contrast, chemorefractory PDOs generally have lower cell-intrinsic mitotic activity than chemosensitive PDOs (Figure 4c). When treated with 5-FU, SN-38, and oxaliplatin, chemorefractory PDOs experience a reduction in S-phase and blocked M-phase consistent with on-target drug responses, but generally elicit a lower double-strand break response compared to chemosensitive patients and do not activate PARP or Caspase3 (Figure 4d) (Figure S7e). This suggests that even chemorefractory PDOs experience on-target drug responses, but their slow mitotic signaling flux at point of treatment means drug-induced cytostatic stress is insufficient to trigger widespread DNA-damage and apoptosis. Chemorefractory PDOs typically have high levels of cell-intrinsic pSMAD2/3, pSMAD1/5/9, pMKK4, pBAD, pBTK, and pNF-κB signaling (Figure 4e) – suggesting these pathways relate to a chemorefactory cell-state. In summary, TOB*is* mass cytometry and Trellis reveal on-target drug performance is common in CRC PDOs (even in chemorefractory PDOs) but cytotoxicity is patient-specific and correlates with cell-intrinsic PDO cell-states and PTM signaling.

### CAFs Chemoprotect PDOs by Altering PDO Cell-State

CAFs have both pro- and anti-cancer roles across a variety of solid tumors, but to what extent these effects are patient-specific is poorly understood [4]. Trellis analysis of all CRC PDOs revealed CAFs can chemoprotect chemosensitive PDOs in a patient-specific manner (Figure 3d). To functionally explore the role of CAFs in patient-specific CRC PDO drug responses, we performed paralleled analysis of PDO mono-cultures and PDO-CAF co-cultures following drug treatments (Figure 5a). Trellis revealed CAFs provide varying degrees of chemoprotection in a patient- and drug-specific manner. For example, CAFs completely protect chemosensitive PDOs 21 and 75 from SN-38, 5-FU, and oxaliplatin-induced apoptosis, whereas PDOs 23, 27, and 99 only experience partial chemoprotection (Figure S10c). Chemorefractory PDOs 05, 11, and 141 are largely unaffected by CAFs. This dichotomy suggests CAFs deregulate cancer cells in a patient-specific manner.

**Figure 5:**
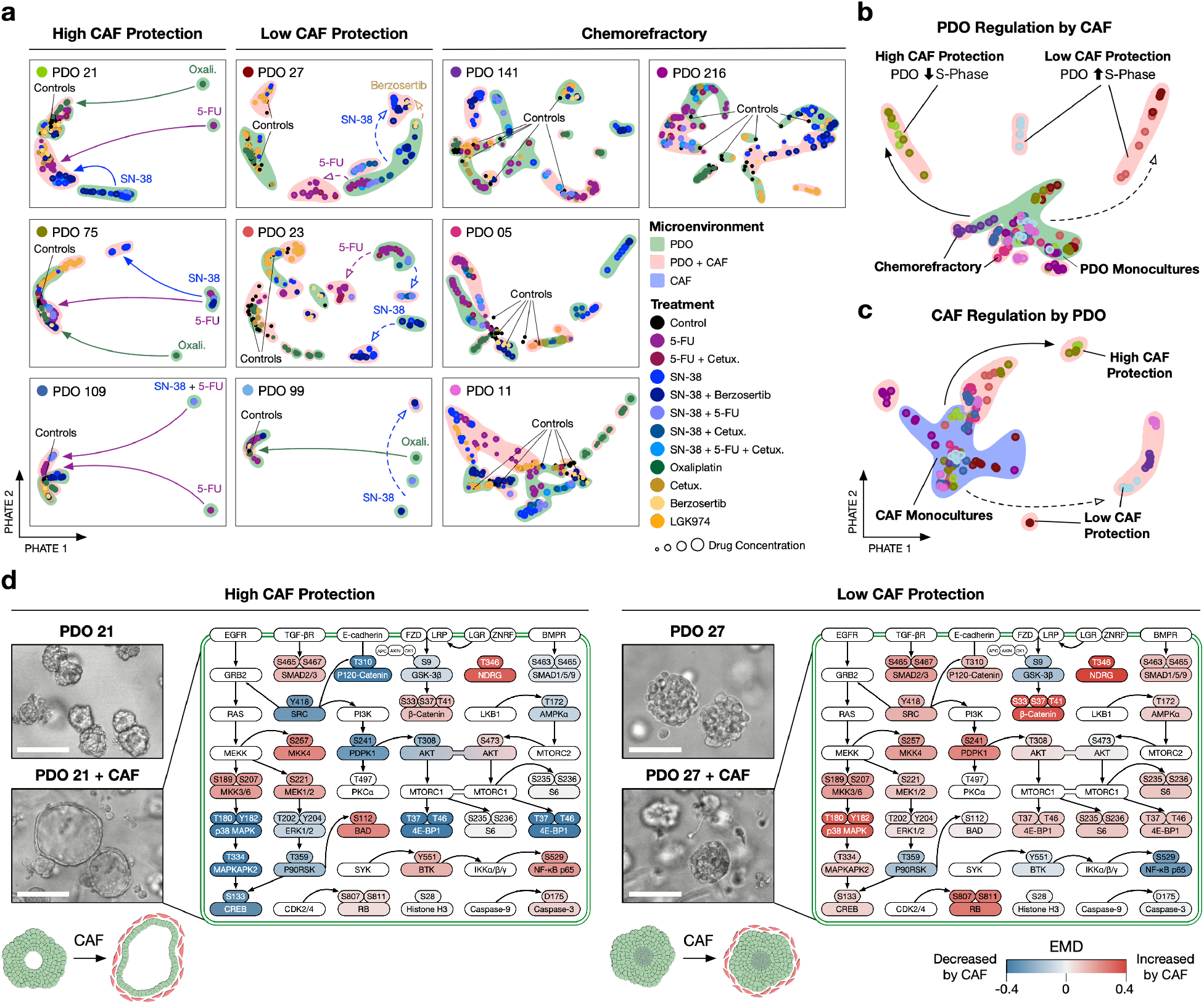
CAFs Chemoprotect PDOs by Altering PDO Cell-State. **a)** Trellis-PHATE of patient-specific PDO PTM drug responses with or without CAFs illustrates CAFs can protect PDOs from therapy (1,680 PDO-CAF cultures). Dots colored by treatment, outlines colored by microenvironment. Solid arrows refer to full protection, dashed arrows refer to low protection by CAFs. **b)** Alterations of PDO cell-state and PTM signaling by CAFs correlates with chemoprotection. Dots correspond to 6 replicates colored by PDO. **c)** Baseline CAF cell-state and PTM signaling when co-cultured with PDOs correlates with chemosensitivity protection. Dots correspond to 6 replicates colored by PDO. **d)** CAF regulation of PTM signaling networks in PDO 21 and PDO 27. CAFs downregulate MAPK and PI3K pathways and upregulate SMAD, NF-*κ*B, and BAD signaling nodes in protected PDOs. Scale bar = 200 *μ*m.

We next sought to understand why CAFs have such different patient-specific regulation of PDO drug response. Chemosensitive PDOs 21 and 75 are highly proliferative in monoculture but reduce cell-cycle activity when co-cultured with CAFs (Figure 5b) (Figure S10a). CAFs that protect PDOs also have a distinct PTM signaling profile in co-culture (Figure 5c), suggesting patient-specific reciprocal signaling between PDOs and CAFs occurs during chemoprotection. Crucially, CAFs do not cause protected PDOs to exit the cell-cycle, but instead reduce MAPK and PI3K signaling, increase TGF-β, JNK, and NF-κB signaling, and slow PDO S-phase entry — rendering PDOs less vulnerable to chemotherapies (Figure 5d). Notably, these pathways are also cell-intrinsically active in chemorefractory PDOs (Figure 4e). CAFs also dramatically alter the macro structure of PDOs, with chemoprotected PDOs switching from an enveloped to cyst-like morphology. PDOs that do not benefit from CAF chemoprotection do not experience morphological shifts. Collectively, we find that CAFs can rapidly regulate PTM signaling networks in PDOs to shift previously chemosensitive cancer cells towards a chemorefractory cell-state.

### Inhibiting YAP Re-sensitizes CAF-Protected PDOs

Mechanistic understanding of drug responses by single-cell signaling analysis could identify opportunities to rationally re-sensitize refractory PDOs [41]. For example, Trellis revealed CAFs protect chemosensitive PDOs from SN-38 — not by reducing on-target S-phase blockage or DNA-damage — but by shifting cancer cells towards a slow-cycling cell-state (Figure 6a-d) (Figure S10a-c). This was most clearly observed in PDO 21, where CAFs activate PDO TGF-β, JNK, and NF-κB signaling and suppress mitotic MAPK and PI3K pathways (Figure 6e). CCD-18Co colon fibroblasts also chemoprotect PDO 21, suggesting PDOs have a common cell-state response to mesenchymal cues (Figure S10e-g).

**Figure 6:**
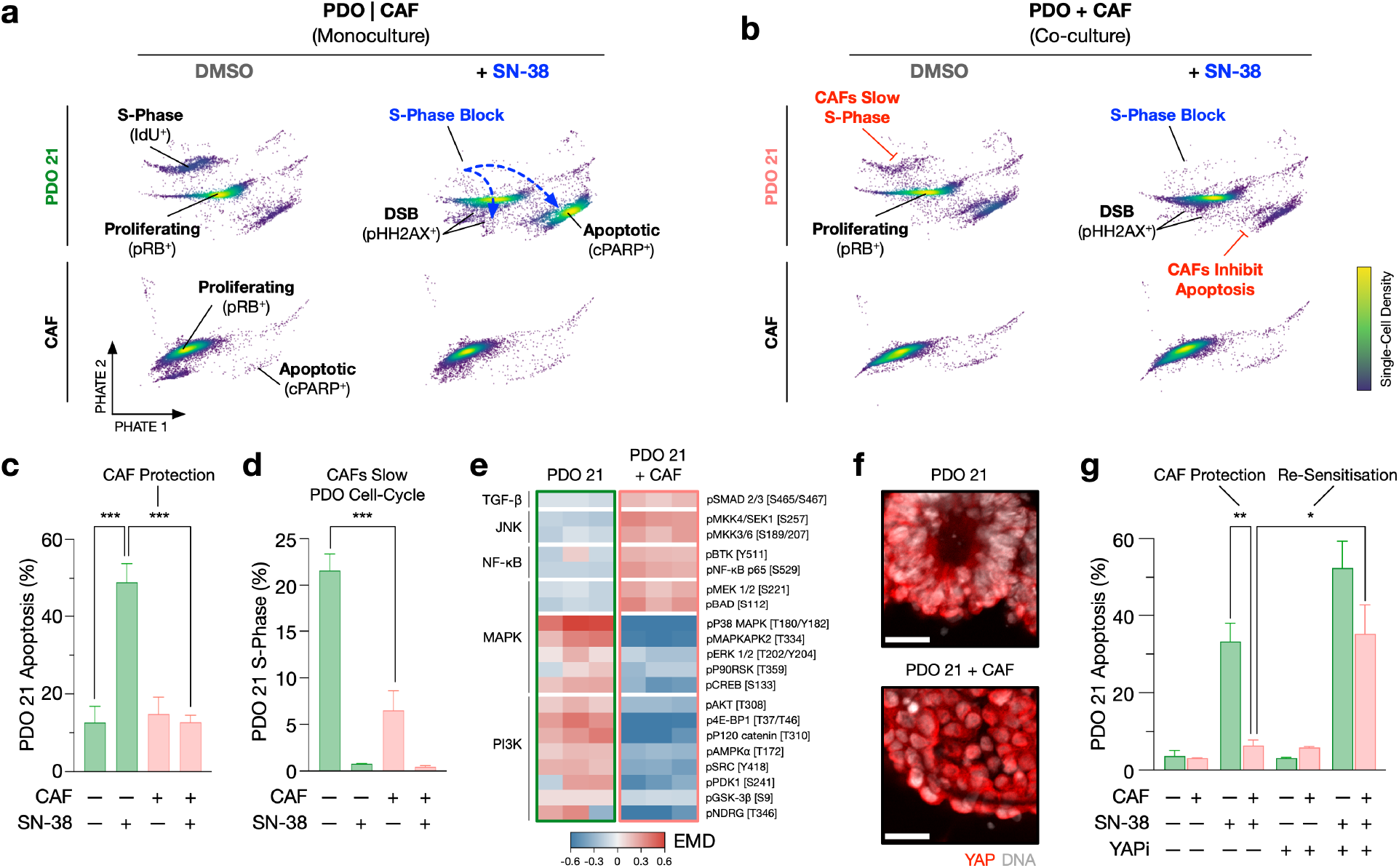
CAF Chemoprotection is Reversed By Inhibiting YAP. **a-b)** Single-cell density PHATEs of PDO 21 and CAFs during SN-38 treatment illustrates cell-state shifts by drug and co-culture (5,000 cells). **c)** CAFs protect PDOs from SN-38-induced apoptosis. **d)** CAFs slow PDO S-phase entry and PDOs experience on-target S-phase blockage by SN-38 irrespective of CAFs. **e)** EMD heatmap of PTMs in PDO 21 +/- CAFs demonstrate CAFs regulate PDO PTM signaling. **f)** CAFs induce nuclear translocation of YAP (red) to PDO nucleus (white). Scale bar = 25 *μ*m **g)** Verteporfin (YAPi) completely re-sensitizes CAF-protected PDOs to SN-38-induced apoptosis. Unpaired *t*-test, *** <0.0001, ** <0.001, * <0.01.

It has recently been shown that CRC cells can escape chemotherapy by differentiating towards a slow-cycling ‘diapause’ [42] or revival stem cell (revSC) fate [43]. In the healthly intestine, revSCs can be induced by fibroblast-derived TGF-β during tissue damage and demonstrate low cell-cycle activity and high levels of SMAD and YAP signaling [44]. While TOB*is* mass cytometry revealed CAF-protected PDOs have low cell-cycle activity, high TGF-β signaling, and low MAPK and PI3K flux, cytometry technologies cannot measure nuclear protein translocation and therefore cannot detect YAP activation. However, YAP immunofluorescence revealed CAFs also induce nuclear YAP translocation in chemoprotected PDOs (Figure 6f) — collectively suggesting CAFs shift PDOs towards a revSC-like cell-state.

Using PTM signaling and cell-state insights provided by single-cell drug screening, we hypothesized that CAF chemoprotection could be YAP-dependent. To test this, we treated PDO 21 + CAF cultures +/- Verteporfin (YAP-TEAD complex inhibitor), +/- SN-38 and measured PTM and cell-state responses using TOBis mass cytometry (Figure 6g). YAP inhibition alone did not induce apoptosis in PDO 21 either in monoculture or in co-culture with CAFs. However, we found Verteporfin completely re-sensitizes CAF-protected PDOs to SN-38-induced apoptosis. Crucially, YAP inhibition did not increase on-target SN-38-induced DNA-damage in PDO 21 and did not regulate CAF cell-cycle or apoptosis. YAP inhibition restored PDOs to an enveloped morphology when in co-culture with CAFs (Figure S11) — indicating YAP inhibition targets the unique CAF-induced PDO cell-state. These results demonstrate that CAFs can chemoprotect PDOs via a YAP-driven revSC cell-state switch and underscore the value of mechanism-focused single-cell drug screening in overcoming therapy resistance.

## 3 Discussion

PDOs have been widely proposed as personalized avatars of patient-specific drug responses [45]. However, bulk screening technologies have limited previous studies to PDO monocultures alone and provide no mechanistic insight into PDO drug response [13]. Using highly-multiplexed single-cell PTM profiling by TOB*is* mass cytometry and hierarchical treatment effect analysis by Trellis, we demonstrate PDO drug response signaling is patient-specific and reveal CAFs regulate PDO chemosensitivity by altering PDO signaling and cell-state. PDO-CAF interactions are also patient-specific, with CAFs both stimulating and repressing PTM signaling and cell-cycle activity in a patient-specific manner. Crucially, we demonstrate mechanistic profiling of patient-specific drug responses can be used to re-sensitize CAF-protected PDOs.

Unlike static diagnostic metrics (e.g. pharmacogenomics) that have failed to substantially advance precision oncology [45], PDOs are functional biopsies that can be experimentally tested to reveal patient-specific drug responses alongside clinical care in real-time [46–48]. However, recent studies have suggested PDOs alone are not sufficient to biomimetically predict drug response. For example, only 20% of monoculture drug combination ‘hits’ could be validated in *ex vivo* organotypic CRC tumours containing a TME [46], and growth factor regulation of PDO cell-state can change pancreatic ductal adenocarcinoma (PDAC) organoid drug responses [49]. Our results reveal PDO-CAF interactions are a source of functional inter-tumor heterogeneity and the role of CAFs should not be generalized. Given that cell-extrinsic signals can have dramatic effects on drug performance, we propose TME cells should be considered in future studies evaluating PDOs as personalized functional biopsies.

Phenotypic plasticity is an emerging hallmark of cancer [50] and therapeutic targeting of cancer-specific cell-states is a growing area of cancer research [51, 52]. As stem cell-driven model systems, PDOs are capable of high differentiation plasticity [8] and are therefore well-suited to studying drug- or TME-induced cancer cell plasticity. We observed that PTM cell-state (not MSI/MSS status, tumor stage, anatomical location, or genotype) aligned with patient-specific drug response (Figure 4c-d) (Figure S9) and found CAFs can transition PDOs into a refractory cell-state to protect PDOs from specific therapies. A recent survey of CRC concluded phenotypic plasticity is largely driven by transcriptional changes, not genotype [53] and work in PDAC has demonstrated PDO transcriptional profiles, not genotype, correlate with drug response [54]. Moreover, recent studies of oncogenic [55] and kinase [56] activity suggest cancer cell signaling flux predicts patient survival better than genotype. Taken with our observations, mounting evidence suggests metrics that more closely describe cancer cell-state such as transcription and PTM signaling may more accurately predict patient-specific drug responses than genomic profiles or clinical staging. Combining the plasticity of PDO models with mechanism-focused single-cell analysis technologies will enable characterization of cell-state plasticity and therapy-induced canalization in cancer.

In contrast to traditional live/dead drug screens, TOB*is* mass cytometry reveals molecular insights into PDO drug responses. We observed PDOs frequently experience on-target drug responses (83%), but only a subset of PDOs enter drug-induced apoptosis (40%). This suggests chemorefractory PDOs do not translate cytostatic and genotoxic stress into apoptosis. Single-cell PTM profiling further revealed CAFs chemoprotect PDOs by shifting cancer cells into a slow-cycling revSC-like cell-state. We used this mechanistic insight to re-sensitize PDOs by blocking revSC activation via YAP. Given that drug synergy is rare when using unbiased screens [57], our study suggests mechanism-focused screening could be used to rapidly identify rational drug synergies to re-sensitize refractory cancers.

The advent of high-dimensional single-cell technologies such as mass cytometry and scRNA-Seq provides new opportunities to study heterogeneous drug response mechanisms beyond simple viability scores [13]. However, high-dimensional drug screening data is challenging to interpret — with existing tools designed to analyze dozens, not thousands of samples. Trellis overcomes this scalability bottleneck by distributing single-cell data across a tree domain structure, enabling the KR norm between thousands of single-cell samples to be computed rapidly. While we use cell-state branches to sensitize Trellis results towards canonical on-target anti-cancer drug responses, alternative branching structures could in theory be designed to enrich for PTM signaling hierarchies (e.g. for kinase inhibitor screens) or cell-type hierarchies (e.g. in immune profiling) (Figure S3). Trellis’ scalability is independent of supervening branches and is therefore a flexible platform for future single-cell screening applications.

Although this study has focused on PDOs and CAFs, single-cell technologies also enable mechanistic analysis of organoid-leukocyte co-culture models [13]. In addition to studying the role of leukocytes in regulating chemical drug responses, single-cell PTM analysis could be a powerful approach to study pre-clinical co-culture organoid models of anti-solid tumor cellular biotherapeutics (e.g. CAR-T cells) where understanding the biology of the drug (engineered T-cell) is as important as the PDO target-cell killing. This study demonstrates high-throughput single-cell screening of heterocellular drug interactions is feasible and we expect the technology will be rapidly adapted to study biological therapies.

In summary, we demonstrate highly-multiplexed single-cell PTM profiling by TOB*is* mass cytometry and hierarchical treatment effect analysis by Trellis can reveal patient-specific drug responses in thousands of PDO-CAF cultures. CAFs regulate PDO drug response by altering PDO cell-state in a patient-specific manner and PTM signaling insights can be used to overcome CAF protection. We propose single-cell PTM analysis as a powerful alternative to traditional bulk viability analysis of PDOs and suggest TME cells should be considered in future precision medicine models.

## 4 Methods

### 4.1 CRC PDO and CRC CAF Culture

CRC PDOs were obtained from the Human Cancer Models Initiative (Sanger Institute, Cambridge, UK) [12] and expanded in 12-well plates (Helena Biosciences 92412T) in x3 25 *μ**L*** droplets of Growth Factor Reduced Matrigel (Corning 354230) per well with 1 mL of Advanced DMEM F/12 (Thermo 12634010) containing 2 mM L-glutamine (Thermo 25030081), 1 mM N-acetyl-L-cysteine (Sigma A9165), 10 mM HEPES (Sigma H3375), 500 nM A83-01 (Generon 04-0014), 10 *μ*M SB202190 (Avantor CAYM10010399-10), and 1X B-27 Supplement (Thermo 17504044), 1X N-2 Supplement (Thermo 17502048), 50 ng ml^-1^ EGF (Thermo PMG8041), 10 nM Gastrin I (Sigma SCP0152), 10 mM Nicotinamide (Sigma N0636), and 1X HyClone Penicillin-Streptomycin Solution (Fisher SV30010), and conditioned media produced as described in [58] at 5 % CO_2_, 37 °C. PDOs were dissociated into single cells with 1X TripLE Express Enzyme (Gibco 12604013) (incubated at 37 °C for 20 minutes) and passaged every 10 days. L-cells for conditioned media production were obtained from Shintaro Sato (Research Institute of Microbial Diseases, Osaka University, Osaka, Japan). To aid cell-type-specific visualization and gating, CRC PDO were transfected with H2B-RFP (Addgene 26001). CRC CAFs (+GFP) were a kind gift from Prof. Olivier De Wever (University of Gent) [16, 17]. CAFs and CCD-18Co fibroblasts (ATCC CRL-1459) were cultured in DMEM (Thermo 11965092) enriched with 10 % FBS (Gibco 10082147), and 1X HyClone Penicillin-Streptomycin Solution (Fisher SV30010) at 5% CO_2_, 37 °C.

### 4.2 PDO-CAF Drug Treatments

PDOs were dissociated into single cells on day 0, and expanded in 12-well plates in Growth Factor Reduced Matrigel (Corning 354230) with Advanced DMEM F/12 (Thermo 12634010) containing 2 mM L-glutamine (Thermo 25030081), 1 mM N-acetyl-L-cysteine (Sigma A9165), 10 mM HEPES (Sigma H3375), 1X B-27 Supplement (Thermo 17504044), 1X N-2 Supplement (Thermo 17502048), 50 ng ml^-1^ EGF (Thermo PMG8041), 10 nM Gastrin I (Sigma SCP0152), 10 mM Nicotinamide (Sigma N0636), 500 nM A83-01 (Generon 04-0014), 10 *μ*M SB202190 (Avantor CAYM10010399-10) and 1X HyClone Penicillin-Streptomycin Solution (Fisher SV30010) at 5% CO_2_, 37 °C for 4 days. On day 5, PDOs were starved in Reduced media (containing only 2 mM L-glutamine, 1 mM N-acetyl-L-cysteine, 10 mM HEPES, 1X B-27 Supplement, 1X N-2 Supplement, 10 mM Nicotinamide, and 1X HyClone-Penicillin Streptomycin Solution) at 5 % CO_2_, 37 °C. In parallel, CAFs were starved in 2 % FBS DMEM with 1X Hyclone-Penincillin Streptomycin Solution. PDOs and CAFs were seeded on day 6 in 96-well plates (Helena Biosciences 92696T) in 50 *μ*L Matrigel stacks with 300 *μ*L of reduced media. PDO monocultures are seeded at a density of ~ 1.5 x10^3^ organoids/well, and CAFs at 2.5 x 10^5^ cells/well, co-cultures were mixed in Matrigel on ice at the densities described, and seeded together on the plates for polymerization. On day 7, media was replaced with titrated concentrations of SN-38 (Sigma H0165), 5-FU (Merck F6627), Oxaliplatin (Merck O9512), Cetuximab (MedChem Express HY-P9905), VX-970 (Stratech), and LGK-974 (Peprotech 1241454) (Table S2) diluted in Reduced media. On day 8, media was replaced with the corresponding treatments (same as on day 7). After 72 hours of co-culture, and 48-hours of treatment (day 9), cultures were processed for TOB*is* mass cytometry (see below). Verteporfin (Cambridge Bioscience CAY17334) was used at 100 nM as above.

### 4.3 PDO-CAF TOBis Mass Cytometry

PDO-CAF co-cultures were analyzed using the TOB*is* mass cytometry protocol outlined in detail by Sufi and Qin *et al*., Nature Protocols, 2021 [15]. Briefly, following drug treatment, PDO-CAF cultures were incubated with 25 *μ*M (5-Iodo-2’-deoxyuridine) (^127^IdU) (Fluidigm 201127) at 37 °C for 30 minutes, and 5 minutes before the end of this incubation, 1X Protease Inhibitor Cocktail (Sigma, P8340) and 1 XPhosSTOP (Sigma 4906845001) are added into the media. After the incubation with ^127^IdU, protease inhibitors and PhosSTOP, each well is fixed in 4 % PFA/PBS (Thermo J19943K2) for 1 hour at 37 °C. PDO-CAFs were washed with PBS, dead cells were stained using 0.25 μM ^194^Cisplatin (Fluidigm 201194), and PDO-CAFs were barcoded *in situ* with 126-plex (9-choose-4) TOBis overnight at 4 °C. Unbound barcodes were quenched in 2 mM GSH and all PDO-CAFs were pooled. PDO-CAFs were dissociated into single-cells using 1 mg ml^-1^ Dispase II (Thermo 17105041), 0.2 mg ml^-1^ Collagenase IV (Thermo 17104019), and 0.2 mg ml^-1^ DNase I (Sigma DN25) in C-Tubes (Miltenyi 130-096-334) via gentleMACS™ Octo Dissociator with Heaters (Miltenyi 130-096-427). Single PDO and CAF cells were washed in cell staining buffer (CSB) (Fluidigm 201068) and stained with extracellular rare-earth metal conjugated antibodies (Table S3) for 30 minutes at room temperature. PDO-CAFs were then permeabilized in 0.1 % (vol/vol) Triton X-100/PBS (Sigma T8787), 50 % methanol/PBS (Fisher 10675112), and stained with intracellular rare-earth metal conjugated antibodies for 30 minutes at room temperature. PDO-CAFs were then washed in CSB and antibodies were cross-linked to cells using 1.6 % (vol/vol) FA/PBS for 10 minutes. PDO-CAFs were incubated in 125 nM ^191^ Ir/^193^Ir DNA intercalator (Fluidigm 201192A) overnight at 4 °C. PDO-CAFs were washed, resuspended in 2 mM EDTA (Sigma 03690) in water (Fluidigm 201069), and analyzed using a Helios Mass Cytometer (Fluidigm) fitted with a ‘Super Sampler’ (Victorian Airships) or CyTOF XT (Fluidigm) at 200-400 events s^-1^.

### 4.4 TOB*is* Mass Cytometry Data Preprocessing

Multiplexed FCS files were debarcoded into separate experimental conditions by using the Zunder Lab Single Cell Debarcoder (https://github.com/zunderlab/single-cell-debarcoder) [19]. Debarcoded FCS files were uploaded to Cytobank (Beckman Coulter), gated for Gaussian parameters, and DNA (^191^Ir/^193^Ir). Epithelial cells were gated on PCK^+^ and EpCAM^+^, and CAFs were gated on Vimentin^+^ and GFP^+^. Arcsinh transformed values were mean centered across batches before Trellis analysis.

### 4.5 Trellis Computational Background

Single-cell data are being collected in experiments with ever more numerous conditions in order to characterize libraries of treatments [59] including small-molecules [60] and gene-perturbations [61]. One method that directly generalizes bulk measurements to single-cell samples is through the theory of optimal transport and more specifically, the Wasserstein distance [22–24].

Optimal transport is well suited to the formulation of distances between collections of points, as it generalizes the notion of distances between points to distances between distributions. Intuitively, the distance between distributions should be the minimum total work to move a pile of dirt located at a source distribution to a target distribution. This framework yields a natural definition of similarity between experimental conditions, namely two conditions are similar when their collections of cells are not far from each other.

These distances aim at answering a deeper question: *Which treatments have similar and different effects on the system?* To answer this question we need a metric between changes to densities. We assume that for each treated condition *X* we have access to an associated control condition *X_c_*. When all treated conditions are measured relative to a single X_c_ we show approaches based on the Wasserstein distance are a valid metric between changes in densities. However, in larger experiments it is impossible to measure all treated conditions within a single batch, and thus treated conditions may have different controls. In this case, we show that Wasserstein-based approaches fail, and show that a generalization to an approach based on the *Kantorovich and Rubinstein norm* gives a valid metric between changes in densities in this more general multi-control case.

#### 4.5.1 Integral Probability Metrics

Integral probability metrics (IPMs) are metrics over probability measures *μ, v* some common space 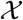 that can be expressed as

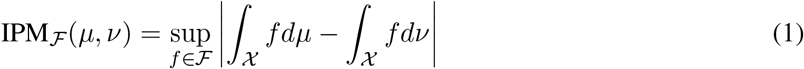

where 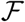 is a family of real-valued bounded measurable functions on 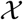. For specific choices of 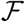 the Dudley metric, Total variation distance, Kologorov distance, maximum mean discrepancy, and Wasserstein distance can all be expressed as IPMs.

IPMs are often useful when we only have samples 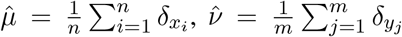 drawn from probability measures *μ, ν* [62]. In this case it is possible to directly estimate IPMs, unlike for the class of *ϕ*-divergences which either do not converge or are +∞. The Wasserstein metric is of particular interest as it has an interpretable primal formulation as the transport of mass between distributions.

#### 4.5.2 The Wasserstein Metric as a Norm

Let *μ, ν* be two probability distributions on a measurable space 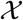 with metric *d*, let Π(*μ, ν*) be the set of joint probability distributions π on the space 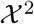 where for any subset 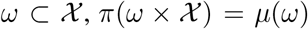 and 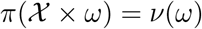. The *α*-Wasserstein distance is defined as:

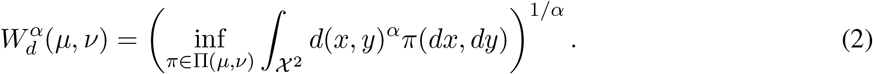

The *Kantorovich–Rubinstein dual* for the Wasserstein distance on arbitrary measures is

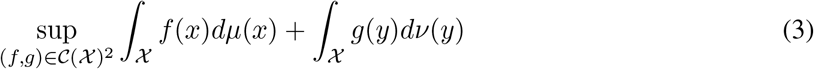

subject to *f*(*x*) + *g*(*y*) ≤ *d*(*x, y*)^*α*^ for all 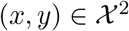. Most work applying the Wasserstein distance focuses on *α* = 2 [63] or more general convex costs with *α* > 1 [64], due to the provable regularity of the transport map. We instead focus on the case where 0 < *α* ≤ 1. Here the transportation map loses regularity but admits a simplification of the dual as when 0 < *α* ≤ 1, it can be shown that Eq. 3 achieves optimality when *g* = –*f* [65, Prop. 6.1] and so simplifies to:

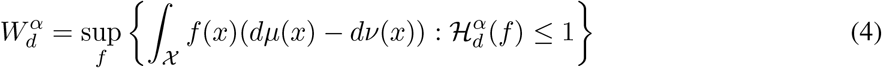

where

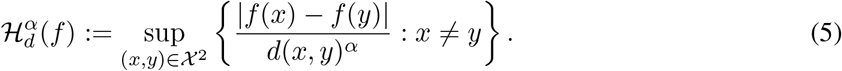

When 0 < *α* ≤ 1, Eq. 4 shows that 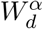 is the dual of the *α*-Hölder functions 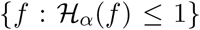 and is a norm, namely

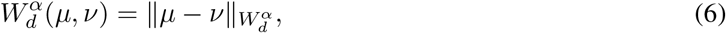

and is valid for any measures *μ, ν* such that 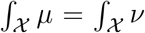. Of particular interest is that 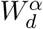 is still a norm even for non-positive measures. This generalization to non-positive measures will form the basis for our Trellis metric between datasets and is known as the *Kantorovich–Rubinstein norm* [66] when applied to differences of non-positive measures.

##### Definition 1

([66]). *The Kantorovich-Rubinstein* (*KR*) *distance between measures μ, ν such that* 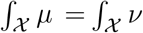 *with respect to ground distance d as*

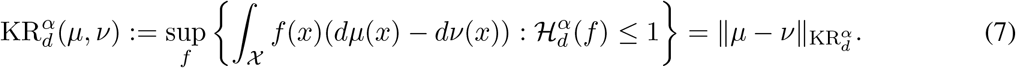

For simplicity we will drop the *α* term and assume *α* = 1, but all statements apply to 0 < *α* ≤ 1 unless otherwise specified. Trellis can be thought of as an efficient implementation of the KR norm over a tree ground distance.

#### 4.5.3 The Wasserstein Distance with Tree Ground Distance

Consider discrete distributions 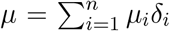 and 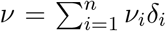 where *δ* is the dirac function in 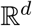 and 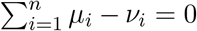. Then for general costs, the Wasserstein distances between *μ* and *ν* can be computed exactly in 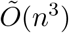 using the Hungarian algorithm [34], and approximated using a slightly modified entropy regularized problem in 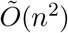 with the Sinkhorn algorithm [35].

However, for some classes of the ground distance, there exist more efficient algorithms (See Table 1). For example, if d is the Euclidean distance in 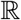, then the Wasserstein distance can be computed in *O*(*n* log *n*) time and is equivalent to sorting [65, 67]. This special case is exploited in sliced-Wasserstein metrics [68, 69] to compute approximate Wasserstein distances in higher dimensions.

**Table 1:** Comparison of Earth Mover’s Distance computation methods separated into super-linear (top), and log-linear methods (bottom) based on time-complexity of computing *k*-Wasserstein-nearest-neighbors. Assumes a dataset of *m* distributions over *n* points with (optionally) a tree of size |*T*| = *O*(*n*).

| Method | Exact | KR-control | Ground cost | knn-Time |
| --- | --- | --- | --- | --- |
| Exact EMD [34] | Yes | No | Any | $O(m^2n^3)$ |
| Sinkhorn EMD [35] | No | No | Any | $O(m^2n^2)$ |
| PhEMD [22] | No | No | $d_{\mathcal{M}}$ | $O(m^2T^3 + n^3)$ |
| Mean | No | Yes | Any | $\tilde{O}(kmn)$ |
| Diffusion EMD [23] | No | Yes | $d_{\mathcal{M}}$ | $\tilde{O}(kmn)$ |
| Trellis / TreEMD (ours) | Yes | Yes | $d_{\mathcal{T}}$ | $\tilde{O}(kmT + n)$ |

Another more general class of ground distances where there exist efficient algorithms is the class of tree metrics. Let 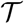 be a rooted tree with non-negative edge lengths, and let 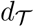 be a *tree metric* on 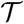. Then for two measures *μ, ν* over 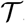, the Wasserstein distance with respect to 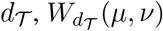, can be computed in *O*(*n*) time by exploiting the fact that there is a single path between any pair of masses [32, 70, 71]. In this case the 1-Wasserstein distance, also known as the Earth Mover’s Distance (EMD) can be expressed as

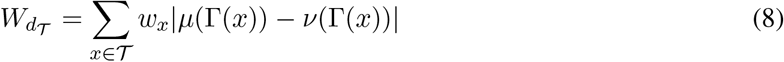

where *w_x_* is the weight / distance to the parent node of *x* and Γ(*x*) represents the set of nodes in the subtree of *x*. Let *P*(*x, y*) be the unique path between *x* and *y*, then 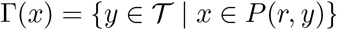. This alternative formulation can be embedded in *l*_1_:

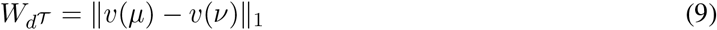

where 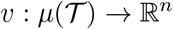 is a function such that *v*(*μ*)_*x*_ = *w_x_μ*(Γ(*x*)).

Approximating the Euclidean distance with a tree distance can be done probabilistically with *O*(*d* log Δ) distortion in expectation where Δ is a resolution parameter [72]. Following the result of Charikar [73], this implies that the 1-Wasserstein distance with tree ground distance has the same order distortion. One simple tree construction that achieves this distortion is known as “Quadtree”, where each node has four children in 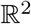 and 2^*d*^ children in 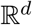 [32]. We introduce a new tree construction based on *k*-means clustering, which we show is a generalization of the Quadtree construction but can be applied to higher dimensions.

##### Algorithm 2: Trellis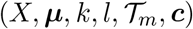

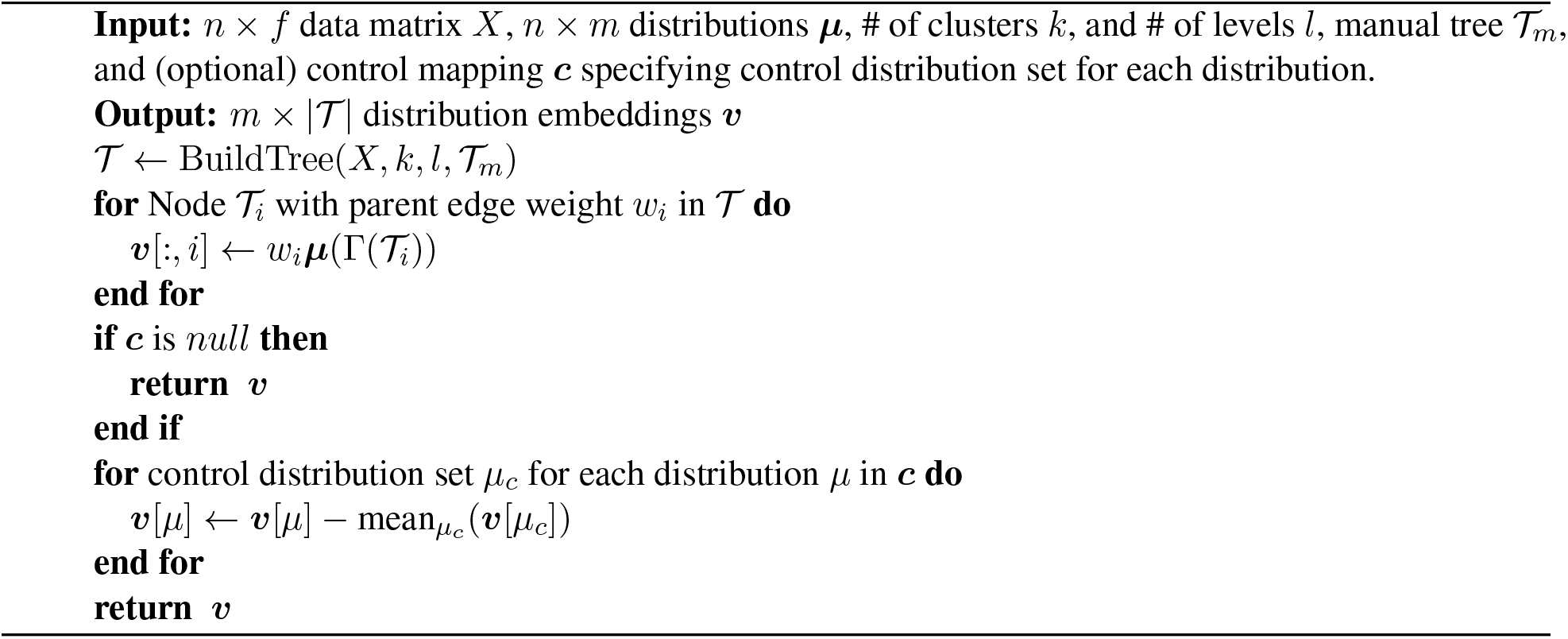

### 4.6 Unpaired and Paired Trellis

We start with a more detailed overview of the Trellis algorithm for comparing the effects of drugs on different experimental conditions. The Trellis algorithm is summarized in Algorithm 2. At a high level Trellis consists of four steps:

1. Construct a hierarchical tree partitioning of the data 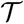.
2. Embed each distribution *μ^i^* over 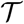 to a vector *v*(*μ^i^*) such that Trellis(*μ^i^, μ^j^*) = ║*ν*(*μ^i^*) – *ν*(*μ^j^*)║_1_ to form a Trellis embedding matrix ***E***.
3. (optionally) Subtract a control distribution embedding 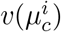 from each *v*(*μ^i^*) for paired Trellis embeddings 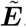.
4. Compute nearest Trellis neighbor distributions exploiting *L*^1^ geometry using fast-nearest-neighbor graph construction algorithms.

We discuss potential methods of constructing 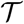 in section 4.6.1, how to embed an empirical distribution to a vector and its equivalence to the Wasserstein distance in section 4.6.2, the effect of subtracting a control distribution embedding in section 4.6.3, and finally how to construct a Trellis-metric nearest neighbor graph for subsequent visualization with a non-linear embedding algorithm such as PHATE [21], UMAP [**?**], or *t*-SNE [28] in section 4.6.4.

#### 4.6.1 Constructing Trees on Single-Cell Mass Cytometry Data

Trellis gives a distance between measures or differences in measures over a tree metric space. Often the data is not associated with an explicit tree metric, but is naturally hierarchical such as in the case of single-cell cytometry data. Previous methods have used manual gating, automatic gating, or a combination of the two to hierarchically cluster single-cell mass cytometry data [74]. These methods build trees, but are missing the ‘metric’ component, which can be encoded as the edge weights between parent and child clusters. We use a simple tree metric where each edge weight for node *x* is the Euclidean distance between the cluster center mean(*x*) and the center of its parent mean(*Pa*(*x*)).

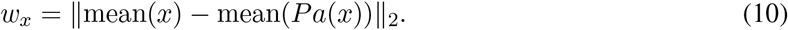

##### Algorithm 3: BuildTree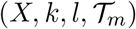

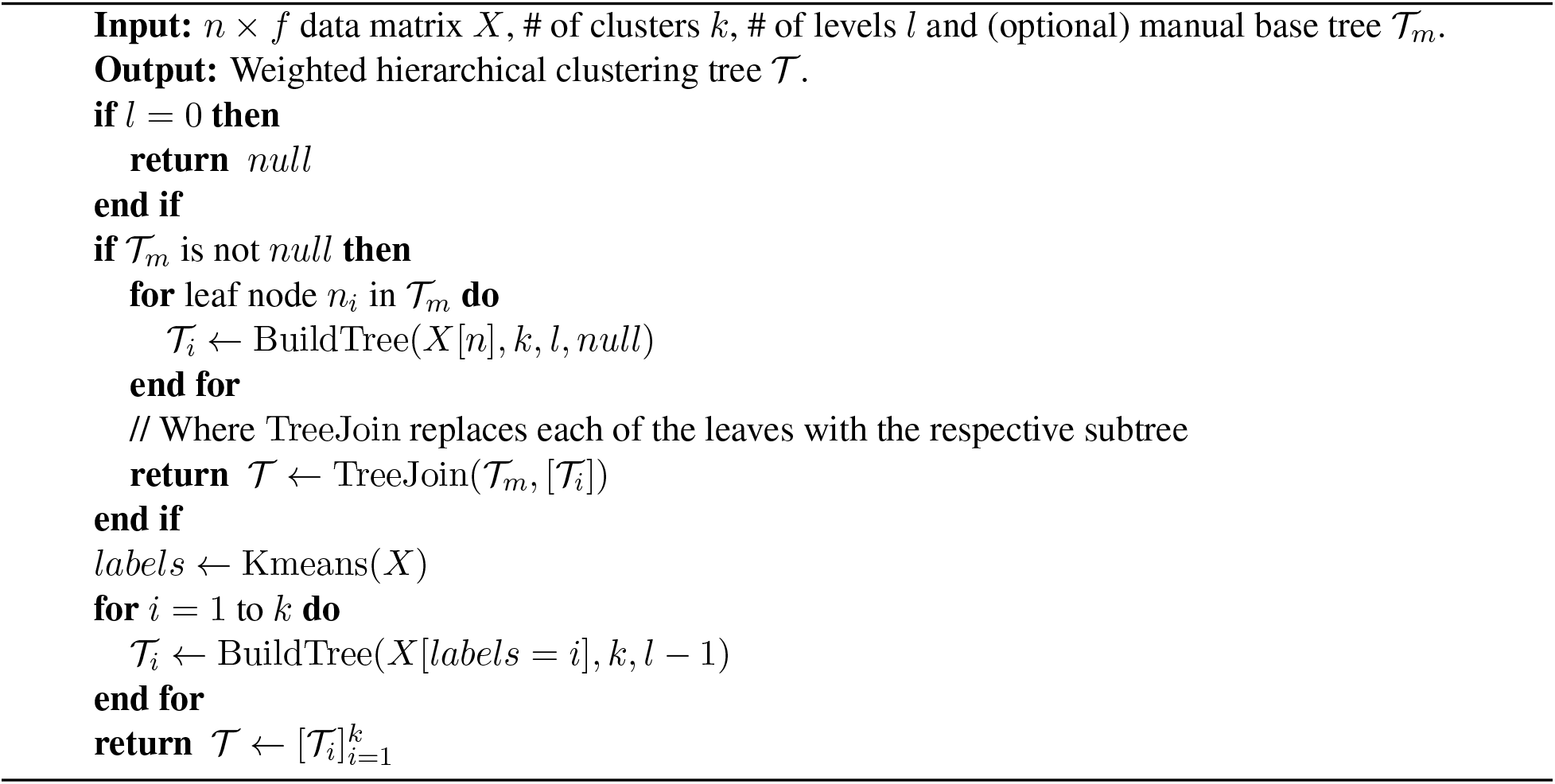

The tree metric between two nodes *u*, 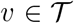 is the sum of the path lengths along the unique path geodesic between *u* and *v* in 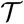 denoted by 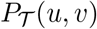 then

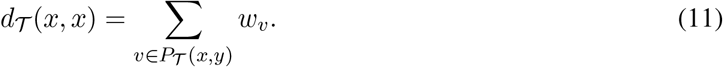

Trellis applies to any clustering method; we demonstrate the Trellis framework using a simple combination of manual gating for non-Euclidean features and automatic gating to approximate Euclidean distances among sub populations. This strategy allows us to leverage manual gating when appropriate due to prior biological knowledge, or automatic gating using repeated *k*-means clustering with no prior on the biological splits. This clustering method is of particular interest because in specific settings we can show that the Trellis metric is topologically equivalent to an Wasserstein distance with Euclidean ground distance in 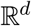.

Given a number of clusters at each level *k* and a depth *h* construct a divisive hierarchical clustering of the data as described in Algorithm 3. Where Kmeans is the *k*-means algorithm with some fixed set of parameters. Interestingly, with a specific setting of *k*-means we show Trellis is topologically equivalent to the α-Wasserstein distance with Euclidean ground distance. This is formalized in the following proposition.

##### Proposition 1.

*Let k* = 2^*d*^, *max_iter* = 0, *data X be normalized such that X* ∈ [–1,1]^*d*^ *with precision* Δ *and initialize the k^th^ cluster at level l with parent center p as p* + 2^1–*l*^ (*Binary*(*k*) – 1/2). *Then there exists constants c, C such that*

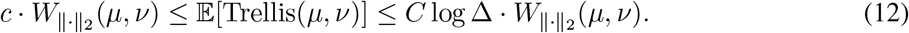

This can be seen by first noting that this initialization is equivalent to a QuadTree construction in the topological sense. If two points are clustered together in our construction at some level then they are also clustered together in QuadTree at the equivalent level. In addition, the edge weights are equivalent up to a constant with the edge weights decaying by 1/2 at every level in both constructions. Once these two properties are verified, then we can leverage existing results on QuadTree constructions from [32] and [73] to show that the inequalities hold. We also note that there exist results on the approximate nearest neighbors of this construction in [71].

While these parameters for kmeans-clustering work well in low dimensions, the number of clusters scales exponentially with dimension. In practice we use four levels of four clusters. This expectation holds over a randomly selected initialization of the zero’th level cluster. In practice, we take the expectation over *k*-means initializations, building ten parallel trees with different initializations.

Trellis can be applied to any tree metric or ensemble of tree metrics. We have presented a method that allows for combining manual and automatic gating, as well as an automatic gating method that in expectation is similar to a Euclidean distance. Many other choices for partitioning CyTof data have been explored in the automatic gating literature [74–77]. These automatic gating methods are generally used for partitioning the data not building a tree metric. However, it is simple to convert them into tree metrics by assigning edge weights based on cluster means. This strategy can be applied to a precomputed hierarchical clustering of the data with no knowledge of how those clusters were chosen. This allows for adaptation of Trellis to different systems where either manual or automatic gating is preferred or already computed.

#### 4.6.2 Trellis Given a Metric Tree

Given a general metric tree 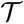 of size 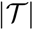, we first define the embedding function 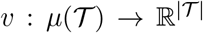 which takes distributions defined over the tree and embeds them in a vector space where the *L*^1^ between vectors is equivalent to the Wasserstein distance with tree ground distance. Given edge weights *w_x_* and denoting the subtree at node *x* as 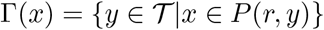, then *ν* is defined element-wise as

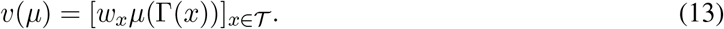

Intuitively, this can be thought of computing the sum of the mass below each node times the edge weight at each node. The difference between *v*(*μ*)_*x*_ – *v*(*v*)_*x*_ for a given node 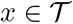 can be thought of as the amount of work needed to move *μ* to *ν*. If this difference is positive, then this means that mass of *μ* is greater in the subtree Γ(*x*) than the mass of *ν*. This means that the transport map must move exactly *μ*(Γ(*x*)) – *ν*(Γ(*x*)) mass upwards from *x* at cost *w_x_*. Adding up these aggregate movements over all nodes gives the total work needed and is equivalent to the work required by the Wasserstein distance.

For our tree construction in Section 4.6.1 with the additional manual tree step, we define the unpaired Trellis distance (uTrellis) as

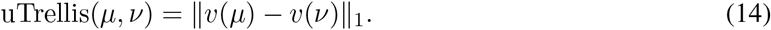

We also define a TreEMD distance without the manual tree construction, considering only the *k*-means construction. TreEMD is similar to previous Tree-based Wasserstein distance constructions for high dimensions [70,71].

These two unpaired distances are comparable to existing methods for computing the Wasserstein distance between distributions. We discuss related methods for computing or approximating the Wasserstein distance in Section 4.6.5. However, these distances do not take into account control, treatment, batch, and replicate information. Given information on which samples were taken under similar conditions, we are able to improve the distances with *Paired Trellis*.

#### 4.6.3 Paired Trellis

To examine the effects of a drug across many conditions it is useful to measure the differences of the treated condition relative to a matched control.

For each sample *μ* and *ν*, let the associated control distributions be *μ_c_* and *ν_c_* respectively, and *v* be defined as above. Then we define the Paired Trellis metric between changes in distributions as:

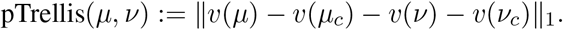

Intuitively, the Paired Trellis distance measures the difference in the change in density between treated conditions from their respective controls. This allows us to control for unmeasured confounders that are implicit in the treated cell population *μ* and *ν* respectively.

##### Proposition 2.

*For two distributions μ, ν with their respective controls μ_c_, ν_c_, the Paired Trellis is equivalent to a Kantorovich-Rubenstein distance with tree ground distance as in Eq. 8*

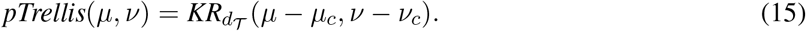

*Proof*. The equivalence of paired Trellis to a Kantorovich-Rubenstein distance can be verified through algebraic manipulation following [78]. We start with the definition of the Kantorovich-Rubenstein distance and show that this is equivalent to pTrellis for an arbitrary tree domain 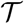 with ground distance 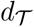. Denote the family of Holder functions under 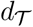 as 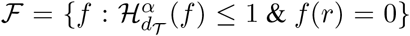 and let λ be the (unique) length measure on 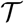 such that 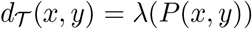. Then there exists a unique function 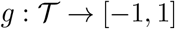 such that: 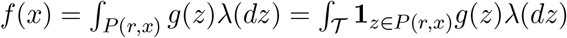.

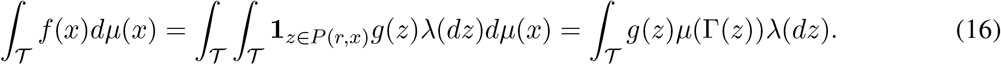

For the optimal witness function *f*^*^, we have

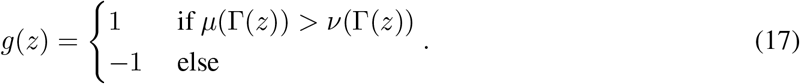

Plugging this equivalence into Eq. 7 we have

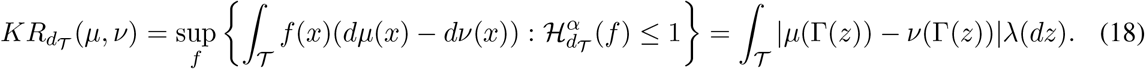

Therefore, for two measures *a, b* over 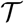 such that 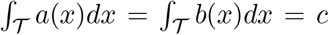 we have that *a*(Γ(*r*)) = *b*(Γ(*r*)) = *c* and for 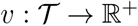 as defined in Eq. 13 we have

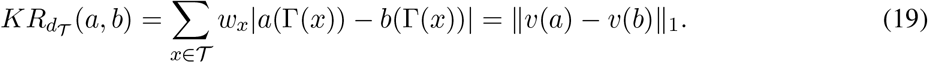

substituting *a* = *μ* – *μ_c_* and *b* = *ν* – *ν_c_* yields the proposition since 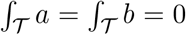 for any distributions *μ, μ_c_, ν*, and *ν_c_*.

We ablate both the pairing and manual tree construction steps in Figure S1. A paired Trellis embedding better separates the effects of increased drug concentration as compared to TreEMD (Figure S1c) and an unpaired Trellis embedding according to a *k*-NN classifier trained with 10-fold cross validation, while also being less sensitive to batch effects by the same metric (Figure S1b).

#### 4.6.4 Nearest Trellis Neighbors

Fast nearest neighbor calculation is useful in graph-based methods which use the k-nearest neighbor graph for down stream tasks such as clustering [79], classification [71], or visualization **[20?**, 21]. For nearest neighbors in normed spaces such as the *L*^2^ norm, the geometry of the space can be utilized for fast exact or approximate nearest neighbor calculation in time scaling logarithmically with the number of points. For more general distances between objects, these algorithms may not apply.

For instance, to compute the *k*-nearest neighbor distributions in terms of the Wasserstein distance for *m* distributions, there is no faster algorithm than computing the Wasserstein distance to all other distributions then computing the *k* closest ones in *O*(*m*) time. However, the Unpaired and Paired Trellis versions of the Wasserstein distance for finite data can be expressed as norms in a finite dimensional space, this allows us to apply fast nearest neighbor algorithms which exploit the induced geometry between distributions. In this case, to find nearest neighbor distributions we can apply tree-based algorithms such as KD-Trees, or Ball-Trees as used in PHATE [21] and scikit-learn [80], locality sensitive hashing in *O*(*T* log *m*) time for m distributions on trees of size *T*.

#### 4.6.5 Related Work and Time Complexity

There are many methods for computing or approximating the Wasserstein distance. In Table 1 we present methods for computing the nearest neighbor distributions according to the Wasserstein distance split into two groups. Here we consider the time it takes for the method to compute the *k*-Wasserstein-nearest-neighbors on a dataset with *m* distributions over *n* points with access to a precomputed tree over the data of size |*T*| = *O*(*n*). The first three methods are widely used, but do not scale well to large datasets with a large number of distributions or a large number of points. For the first three methods, the Hungarian algorithm [34], the Sinkhorn algorithm [35], and PhEMD [22], to find the *k*-nearest-neighbors for a distribution it is necessary to compute the distance to all *m* other distributions. This implies that they scale poorly with the number of distributions as illustrated in Figure S2b. PhEMD saves significant time by only computing the distances between a small set of clusters, however, eventually this is dominated by an increasing number of distributions.

Trellis and TreEMD scale log linearly in the number of points, distributions, and the size of the precomputed tree 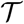. Constructing the tree partitioning for Trellis takes *Õ*(*n*) time. Embedding the distributions takes *O*(*mT*) time. Subtracting the control distribution embedding for paired Trellis takes *O*(*T*) time. finally, computing the *k*-nearest neighbors of the Trellis distance takes *Õ*(*kmT*) time. In total both unpaired and paired Trellis take *Õ*(*kmT* + *n*) time to compute the k nearest neighbor distributions.

When *T* ≪ *n* as in our case, we can see substantial increases in speed in line with simply taking the Euclidean distance between means of clusters. As *T* achieves its upper bound of 2*n* – 1, Trellis has the same complexity as computing the nearest distribution means and of DiffusionEMD [23].

## 5 Data Availability

All mass cytometry files are available on Cytobank at: https://community.cytobank.org/cytobank/projects/1461 Compiled TOB*is* mass cytometry PDO-CAF dataframe is available at: https://data.mendeley.com/datasets/hc8gxwks3p (with a key in Table S2).

## 6 Code Availability

Trellis code is available at: https://github.com/KrishnaswamyLab/Trellis. Code to reproduce all PHATE embeddings in this paper is available at: https://github.com/TAPE-Lab/Ramos-et-al-Trellis.

## 7 Acknowledgments

We are extremely grateful to M. Garnett, H. Francies and the Cell Model Network UK for sharing CRC PDOs and O. De Wever for providing CRC CAFs. We thank Y. Guo, K. Boustani, and G. Morrow from the UCL CI Flow-Core for mass cytometry support. This work was supported by Cancer Research UK (C60693 / A23783), the Cancer Research UK City of London Centre (C7893 / A26233), the UCLH Biomedical Research Centre (BRC422), the Rosetrees Trust (M872 / A2292), the Yale-UCL Collaborative Student Exchange Programme, the NIH (R01GM135929 / R01GM130847), the NSF Career (2047856), the Chan-Zuckerberg Initiative (CZF2019-182702 / CZF2019-002440), and the Sloan Fellowship (FG-2021-15883).

## 8 Author Contributions

M.R.Z. designed the study, performed all PDO-CAF TOBis mass cytometry experiments, analyzed the data, and wrote the paper. A.T. conceived and developed Trellis, analyzed the data, and wrote the paper. J.S. conjugated mass cytometry antibodies and developed TOBis barcodes. P.V., C.N., and X.Q. provided PDO and CAF support. F.C.R. analyzed the data. D.H. provided chemotherapies and oversaw the project. S.K. conceived and oversaw Trellis. C.J.T. designed the study, analyzed the data, and wrote the paper.

---

**Table S1:**
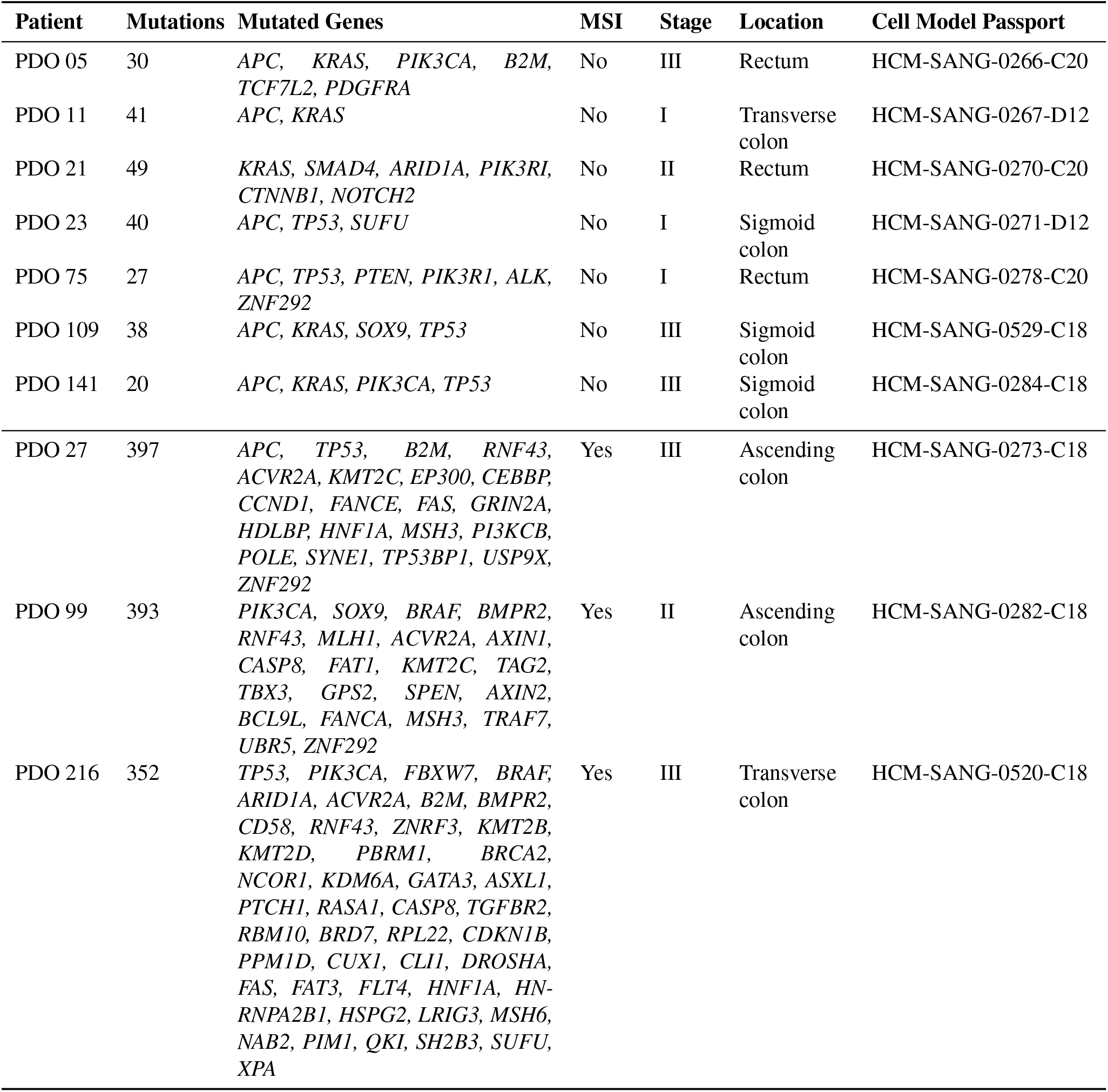
PDO mutations and clinical metadata.

**Table S2:** Drug treatment key for PDO-CAF TOBis mass cytometry master dataframe (Ramos, Maria (2022), “Ramos Zapatero et. al (Cancer-Associated Fibroblasts Regulate Patient-Derived Organoid Drug Responses)”, Mendeley Data, V1, doi: 10.17632/hc8gxwks3p.1).

| Treatment | Label | Conc. Label | Conc. (1) | Conc. (2) | Conc. (3) |
| --- | --- | --- | --- | --- | --- |
| SN-38 | S | 1 | 1 nM | — | — |
|  | S | 2 | 10 nM | — | — |
|  | S | 3 | 50 nM | — | — |
|  | S | 4 | 100 nM | — | — |
| 5-FU | F | 1 | 200 nM | — | — |
| | F | 2 | 2 μM | — | — |
| | F | 3 | 20 μM | — | — |
| | F | 4 | 200 μM | — | — |
| Oxaliplatin | O | 1 | 2 nM | — | — |
|  | O | 2 | 20 nM | — | — |
|  | O | 3 | 200 nM | — | — |
| LGK-974 | L | 1 | 1 nM | — | — |
|  | L | 2 | 10 nM | — | — |
|  | L | 3 | 50 nM | — | — |
|  | L | 4 | 5000 nM | — | — |
| SN-38 (1) + VX-970 (2) | V | 1 | — | 250 nM | — |
|  | VS | 2 | 1 nM | 250 nM | — |
|  | VS | 3 | 10 nM | 250 nM | — |
|  | VS | 4 | 100 nM | 250 nM | — |
| SN-38 (1) + 5-FU (2) + Cetux. (3) | C | 1 | — | — | 5 nM |
|  | CS | 2 | 10 nM | — | 5 nM |
| | CF | 3 | — | 2 μM | 5 nM |
| | SF | 4 | 10 nM | 2 μM | — |
| | CSF | 5 | 10 nM | 2 μM | 5 nM |
| DMSO | DMSO | 0 | — | — | — |
| NH<sub>4</sub>OH | AH | 0 | — | — | — |
| H<sub>2</sub>O | H<sub>2</sub>O | 0 | — | — | — |

**Table S3:** Mass cytometry antibody panel used in all TOBis PDO-CAF experiments.

| Extracellular Antigen | Metal | Clone | Supplier |
| --- | --- | --- | --- |
| CD326 (EpCAM) | <sup>113</sup>In | 9C4 | BioLegend |
| CHGA | <sup>170</sup>Er | C-12 | Insight Biotechnology |
| CD90 (THY1) | <sup>173</sup>Yb | 5E10 | CST |
| Intracellular Antigen | Metal | Clone | Supplier |
| pHistone H3 [S28] | <sup>89</sup>Y | HTA28 | BioLegend |
| RFP | <sup>106</sup>Cd | 8E5.G7 | eBiosciences |
| mCherry | <sup>110</sup>Cd | 16D7 | ThermoFisher |
| Vimentin | <sup>111</sup>Cd | RV202 | BD Biosciences |
| CK18 | <sup>114</sup>Cd | C-04 | Abcam |
| Pan-CK | <sup>115</sup>In | AE1/AE3 | Biolegend UK |
| GFP | <sup>116</sup>Cd | 5F12.4 | eBiosciences |
| pPDK1 [S241] | <sup>141</sup>Pr | J66-653.44.22 | BD Biosciences |
| cCaspase 3 [D175] | <sup>142</sup>Nd | D3E9 | CST |
| Geminin | <sup>143</sup>Nd | Polyclonal | Santa Cruz |
| pMEK 1/2 [S221] | <sup>144</sup>Nd | 166F8 | CST |
| pNDRG1 [T346] | <sup>145</sup>Nd | D98G11 | CST |
| pMKK4 SEK1 [S257] | <sup>146</sup>Nd | C36C11 | CST |
| pBTK [Y511] | <sup>147</sup>Sm | 24a/BTK | BD Biosciences |
| pSRC [Y418] | <sup>148</sup>Nd | SC1T2M3 | BD Biosciences |
| p4E-BP1 [T37/46] | <sup>149</sup>Sm | 236B4 | CST |
| pRB [S807/811] | <sup>150</sup>Nd | J1112-906 | BD Biosciences |
| pPKCα [T497] | <sup>151</sup>Eu | K14-984 | BD Biosciences |
| pAKT [T308] | <sup>152</sup>Sm | J1-223.371 | BD Biosciences |
| pCREB [S133] | <sup>153</sup>Eu | 87G3 | CST |
| pSMAD1/5/9 [S463/465] / [S463/465] / [S465/467] | <sup>154</sup>Sm | D5B10 | CST |
| pAKT [S473] | <sup>155</sup>Gd | D9E | CST |
| pNK-κB p65 [S529] | <sup>156</sup>Gd | K10-895.12.50 | BD Biosciences |
| pMKK3/pMKK6 [S189/207] | <sup>157</sup>Gd | D8E9 | CST |
| pP38 [T180/Y182] | <sup>158</sup>Gd | D3F9 | CST |
| pMAPKAPK2 [T334] | <sup>159</sup>Tb | 27B7 | Abcam |
| pAMPKα [T172] | <sup>160</sup>Gd | 40H9 | CST |
| pBAD [S112] | <sup>161</sup>Dy | 40A9 | CST |
| pHistone H2A.X [S139] | <sup>162</sup>Dy | D7T2V | CST |
| pP90RSK [T359] | <sup>163</sup>Dy | D1E9 | CST |
| p120 Catenin [T310] | <sup>164</sup>Dy | 22/p120 (pT310) | BD Biosciences |
| β-Catenin [Active] | <sup>165</sup>Ho | D13A1 | CST |
| pGSK-3β [S9] | <sup>166</sup>Er | D85E12 | CST |
| pERK1/2 [T202/Y204] | <sup>167</sup>Er | 20A | BD Biosciences |
| pSMAD2/3 [S465/467] / [S423/425] | <sup>168</sup>Er | D27F4 | CST |
| PLK1 | <sup>169</sup>Tm | 35-206 | ThermoFisher |
| pDNAPK [S2056] | <sup>171</sup>Yb | EPR5670 | Abcam |
| pS6 [S235/236] | <sup>172</sup>Yb | D57.2.2E | CST |
| cPARP [D214] | <sup>174</sup>Yb | D64E10 | CST |
| pCHK1 [S345] | <sup>175</sup>Lu | 133D3 | CST |
| Cyclin B1 | <sup>176</sup>Yb | GNS-11 | BD Biosciences |

**Figure S1:**
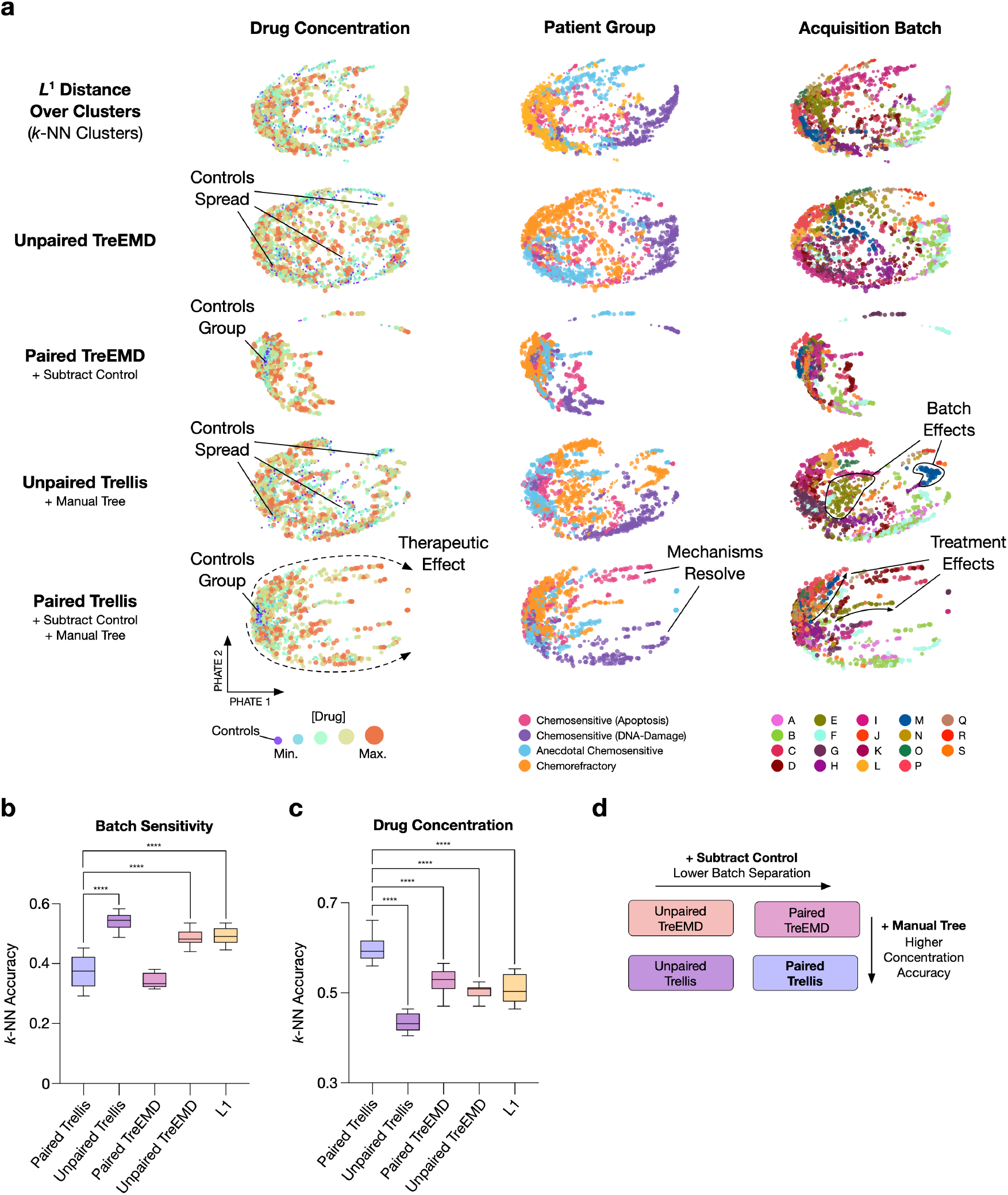
Trellis Ablation Test. **a)** Comparison of Trellis’ ablated algorithm into: L^1^ distance over *k*-NN clusters, Wassertein distance over automatic gating (Unpaired TreEMD), Kantorovich-Rubenstein (KR) norm over automatic gating (Paired TreEMD), Wassertein distance over hierarchical tree partitions of the data by cell-state (Unpaired Trellis), and KR norm hierarchical tree partitions of the data by cell-state (Paired Trellis). **b)** *k*-NN accuracy score on acquisition batches. A higher k-NN accuracy infers a higher batch separation effect by the method. **c)** *k*-NN accuracy score on drug concentrations vs controls. Paired Trellis improves drug treatment effect detection. **d)** Schematic representation of the comparison across methods. One-way ANOVA, **** = <0.0001 (n=10).

**Figure S2:**
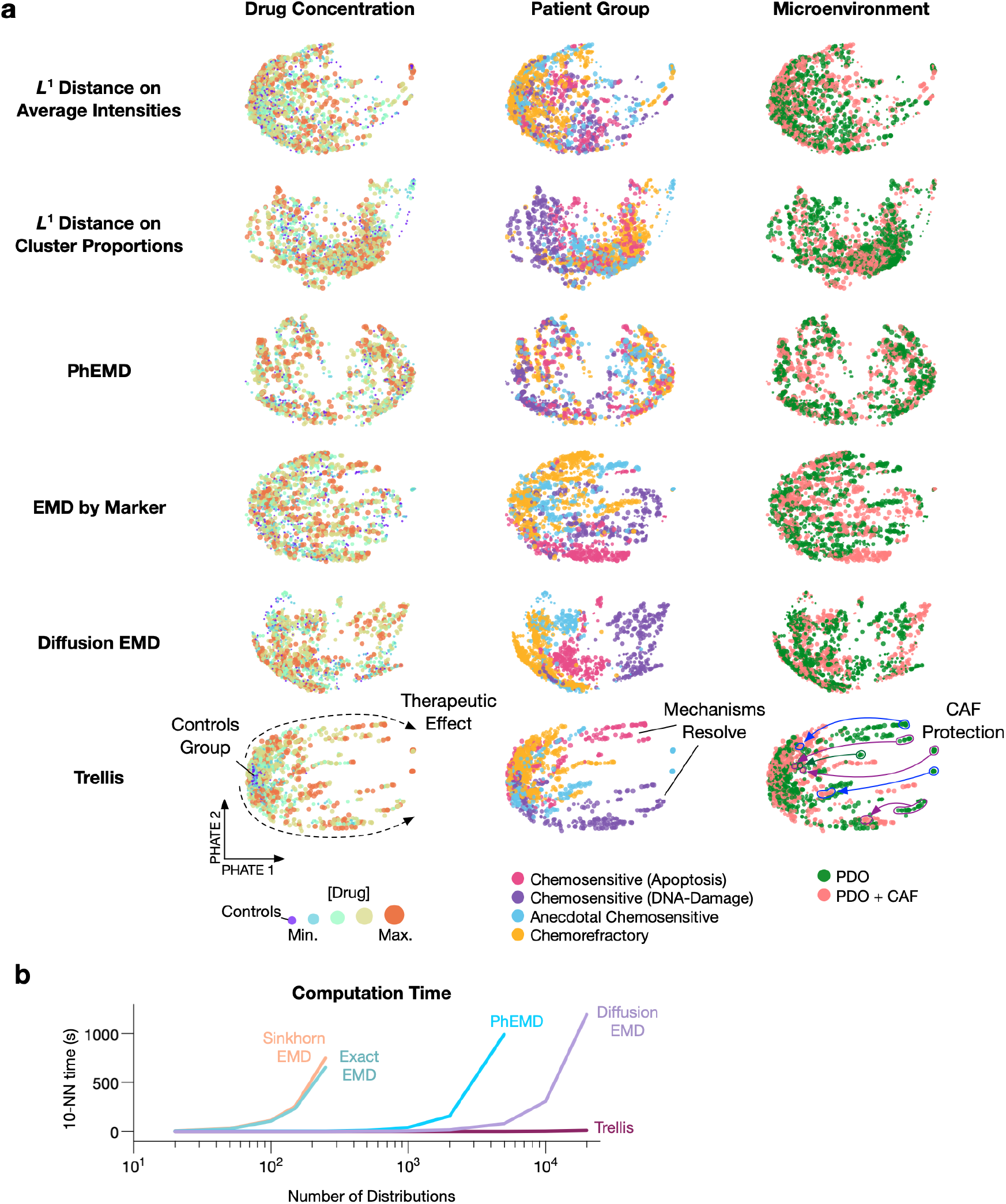
Comparison of Trellis to Alternative Methods. **a)** Trellis performance compared to existing methods such as *L*^1^ distance of average intensity of the markers, *L*^1^ distance of differential abundance of cells in clusters, PhEMD, EMDs between samples on marker intensities, and Diffusion EMD. Alternative methods fail to capture therapeutic effects and cannot identify CAF protection. **b)** Trellis speed and scalability relative to alternative EMD methods.

**Figure S3:**
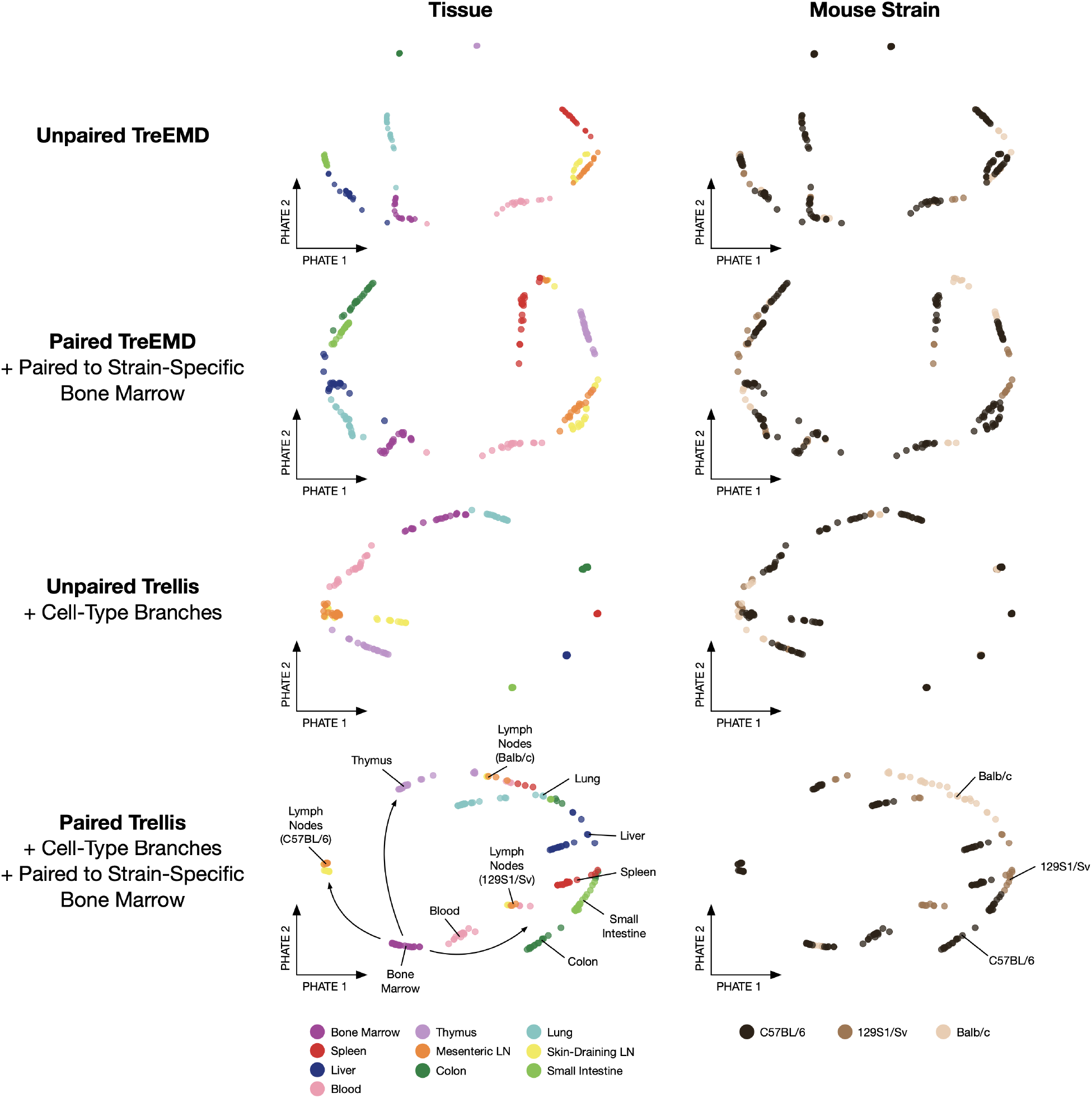
Trellis Analysis of Murine Immune Cell Atlas. Unpaired TreEMD, Paired TreEMD (paired to bone marrow control), Unpaired Trellis (using immune cell-type branches), and Paired Trellis (using immune cell-type branches, paired to bone marrow control) analysis of murine immune atlas mass cytometry data (from Spitzer *et al*., Science, 2015 [81]) (202 single-cell datasets). All tree-based methods resolve tissue-specific immune profiles, but Paired Trellis also captures broad haematopoietic development trajectories and reveals mouse strain specific differences (specifically regarding strain-specific lymph node profiles).

**Figure S4:**
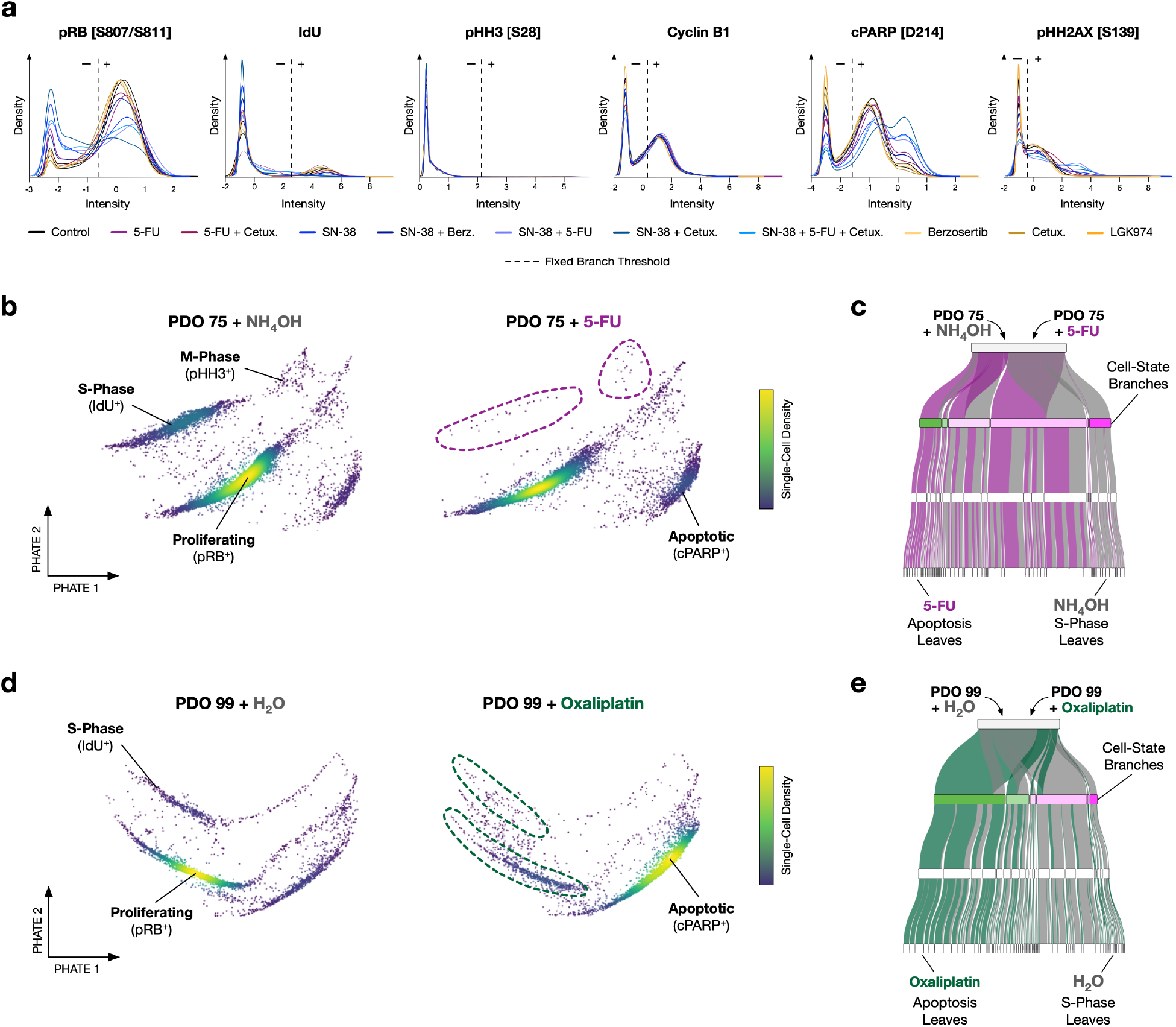
Trellis Detection of PDO Cell-State Drug Responses. **a)** Trellis cell-state branch thresholds for PDO 21 (batch-mean centered and arcsinh transformed intensities). **b)** Single-cell density PHATEs of PDO 75 treated with NH_4_OH vehicle control or 5-FU. **c)** Sankey diagram showing data from b) distributing through the cell-state Trellis layout in Fig. 2 (terminal leaves not shown). **d)** PDO 99 treated with H_2_O vehicle control or Oxaliplatin. **e)** Sankey diagram showing data from d) distributing through the cell-state Trellis layout in Fig. 2 (terminal leaves not shown).

**Figure S5:**
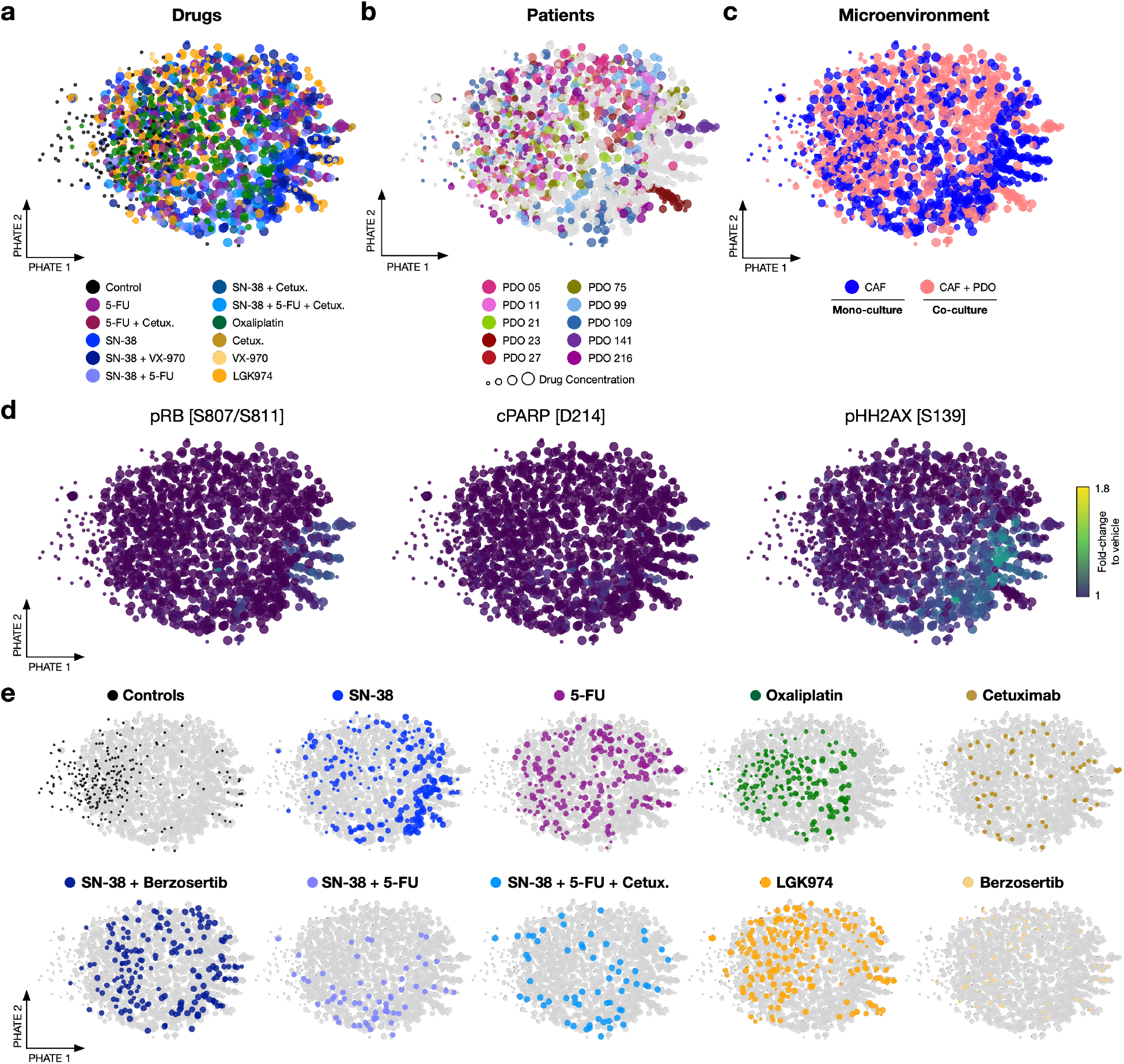
Single-Cell PTM CAF Drug Responses. **a)** Trellis-PHATE of PTM profiles from PDO-CAF cultures fails to identify drug-specific CAF responses, **b)** patient-specific CAF drug responses, or **c)** microenvironment-specific CAF drug responses. **d)** Fold-change to vehicle of pRB [S807/S811], cPARP [D214], and pHH2AX [S139] fail to resolve drug- or patient-specific shifts in cell-state. **e)** CAF responses to individual drug treatments.

**Figure S6:**
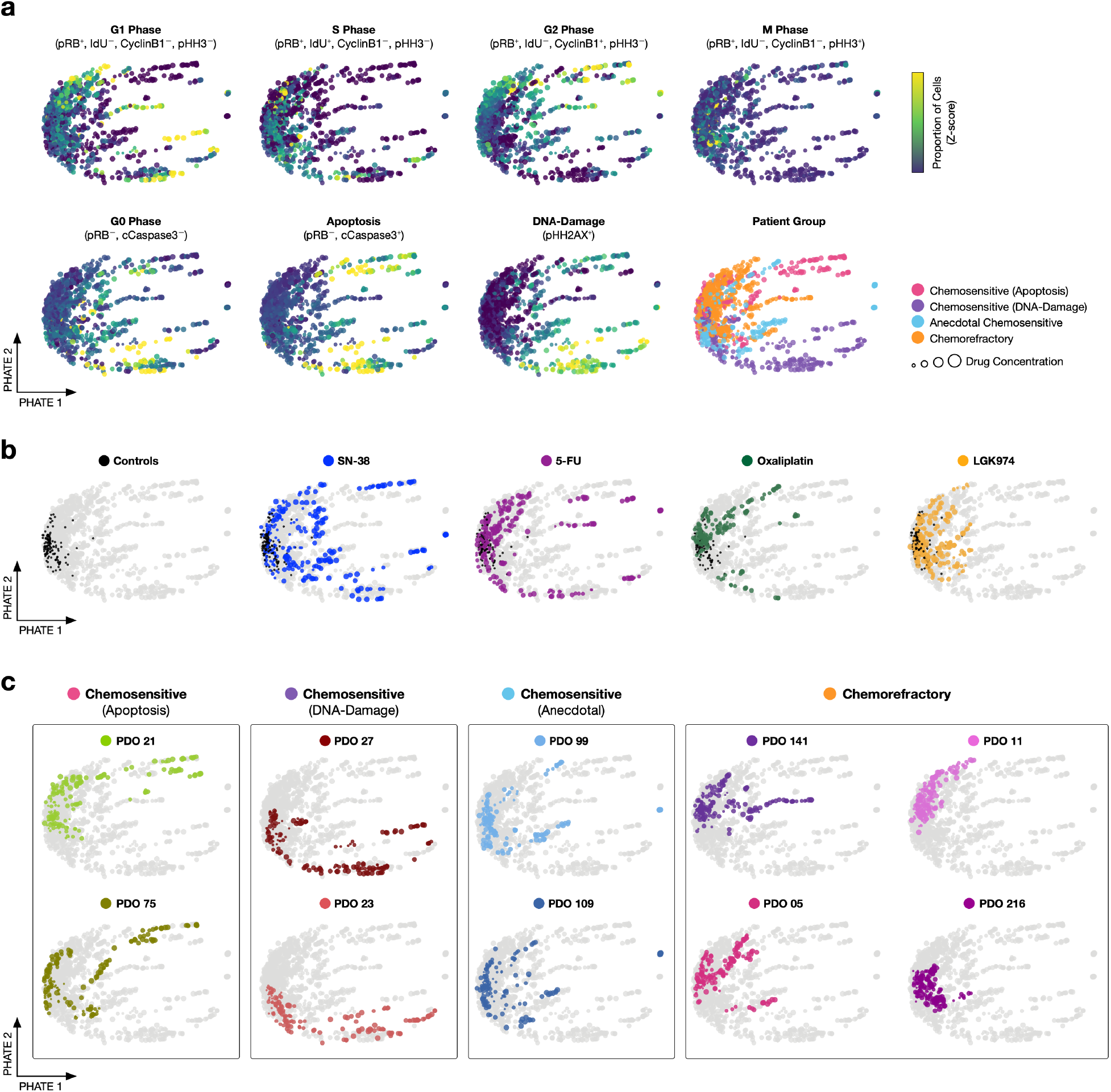
PDO Trellis-PHATE Cell-State Distributions. **a)** Treatment cell-state (z-score) across 1,680 single-cell PDO cultures reveal mechanistic drug treatment effects. **b)** Control, SN-38, 5-FU, Oxaliplatin, and LGK974 distributions. **c)** Individual patient distributions.

**Figure S7:**
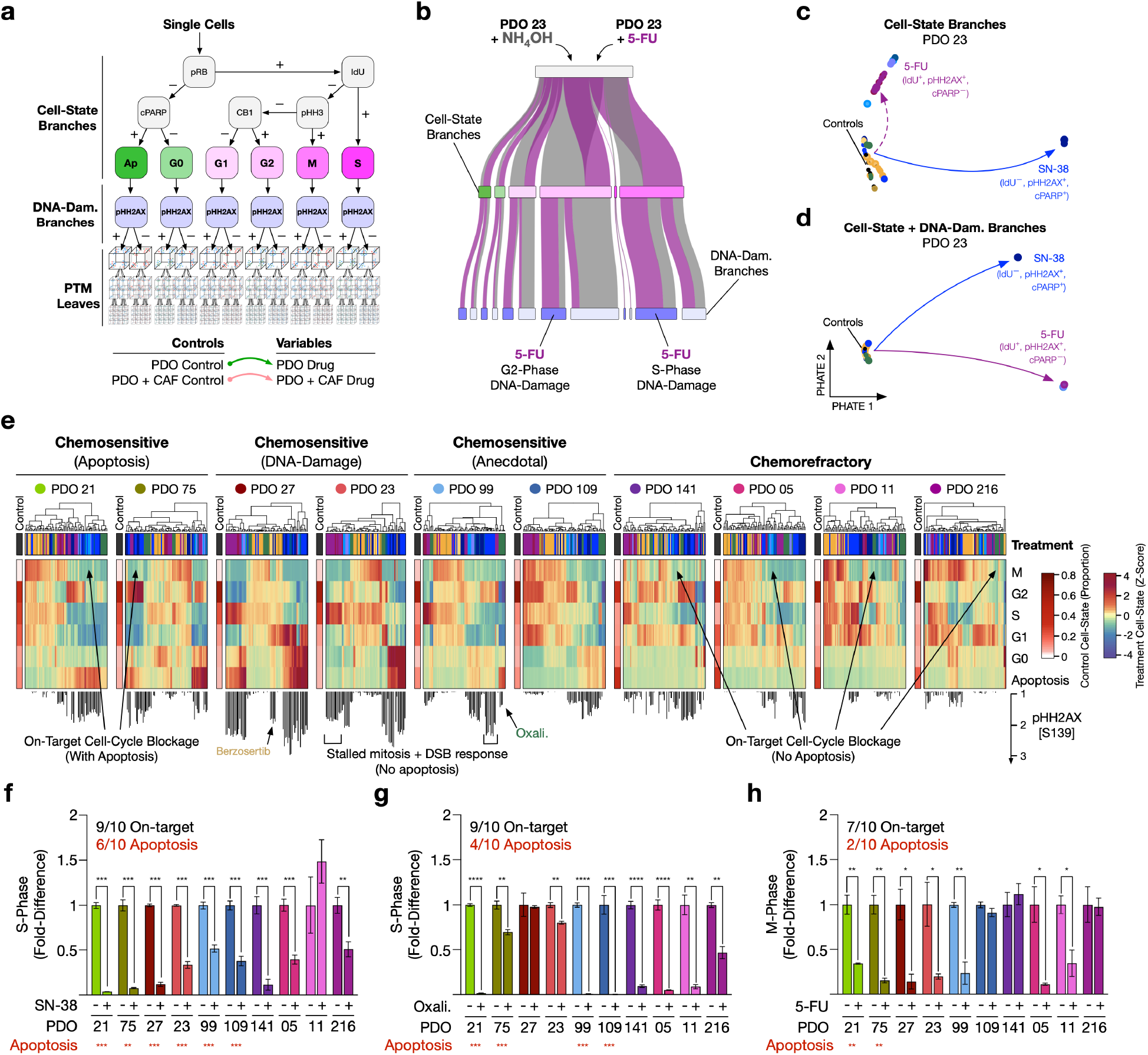
Patient-Specific Regulation of Cell-State and DNA-Damage. **a)** Trellis hierarchy containing cell-state branches with a pHH2AX [S139] DNA double-strand break detection layer. **b)** Sankey diagram showing NH4OH vehicle control and 5-FU treatment of PDO 23 distributing through the cell-state Trellis branches in a) (leaves not shown). **c)** Trellis-PHATE of PDO 23 treatments analyzed using cell-state branches alone or **d)** cell-state branches and pHH2AX [S139] detection layer. The DNA-damage detection layer improves resolution of 5-FU on-target treatment effect. Solid arrows refer to strong treatment effect, dashed arrows refer to partial treatment effect. **e)** Patient-specific distribution of cells within Trellis branches reveals mechanistic cell-state shifts and DNA-damage upon drug treatments. Treatment cell-state quantifies the fold change of the proportion of cells/cell state over the controls for each treatment (Z-score). DNA damage is quantified by the fold change of the proportion pHH2AX^+^ cells over the controls. **f)** PDO cells in S-phase following 100 nM SN-38. **g)** PDO cells in S-phase following 200 nM Oxaliplatin. **h)** PDO cells in M-phase following 200 nM 5-FU. PDOs with a significant >1.5 fold increase in apoptosis are indicated in red. Unpaired t-test, *** <0.0001, ** <0.001, * <0.01.

**Figure S8:**
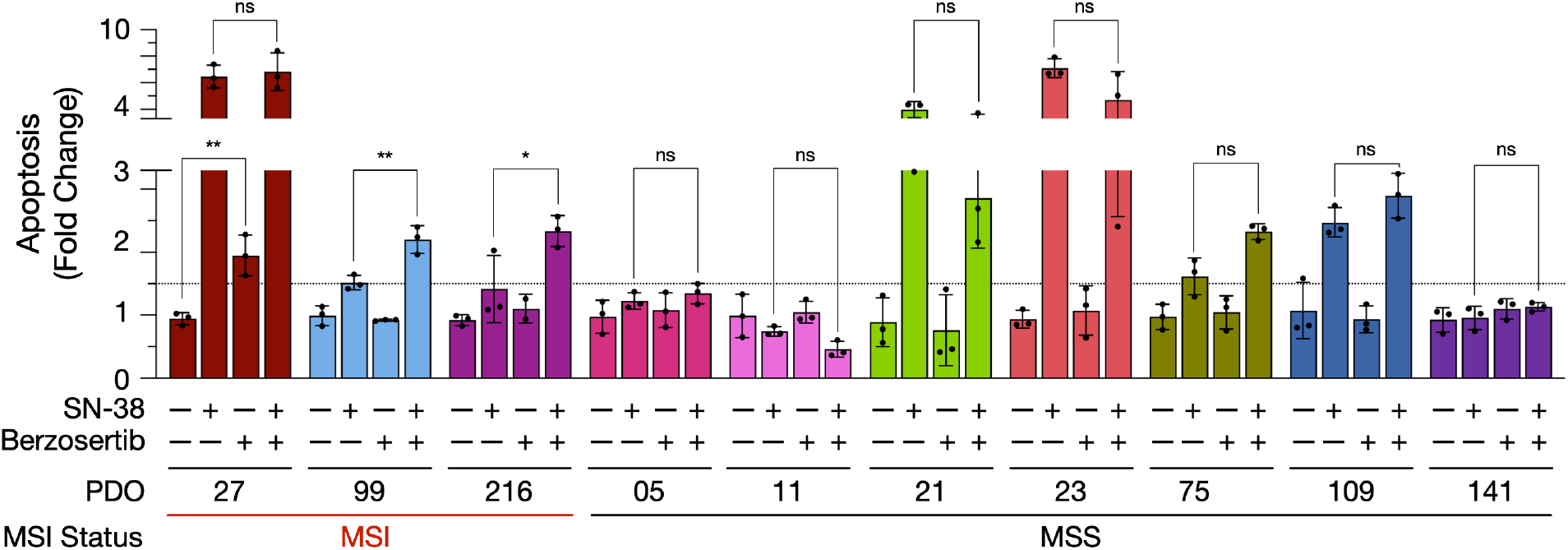
ATR Inhibitor Sensitivity. PDO apoptosis following treatment with SN-38 and/or Berzosertib. Only MSI PDOs are sensitive to ATR inhibitors either alone (PDO 27) or in combination with SN-38 (PDOs 99 and 216). Unpaired t-test, *** <0.0001, ** <0.001, * <0.01.

**Figure S9:**
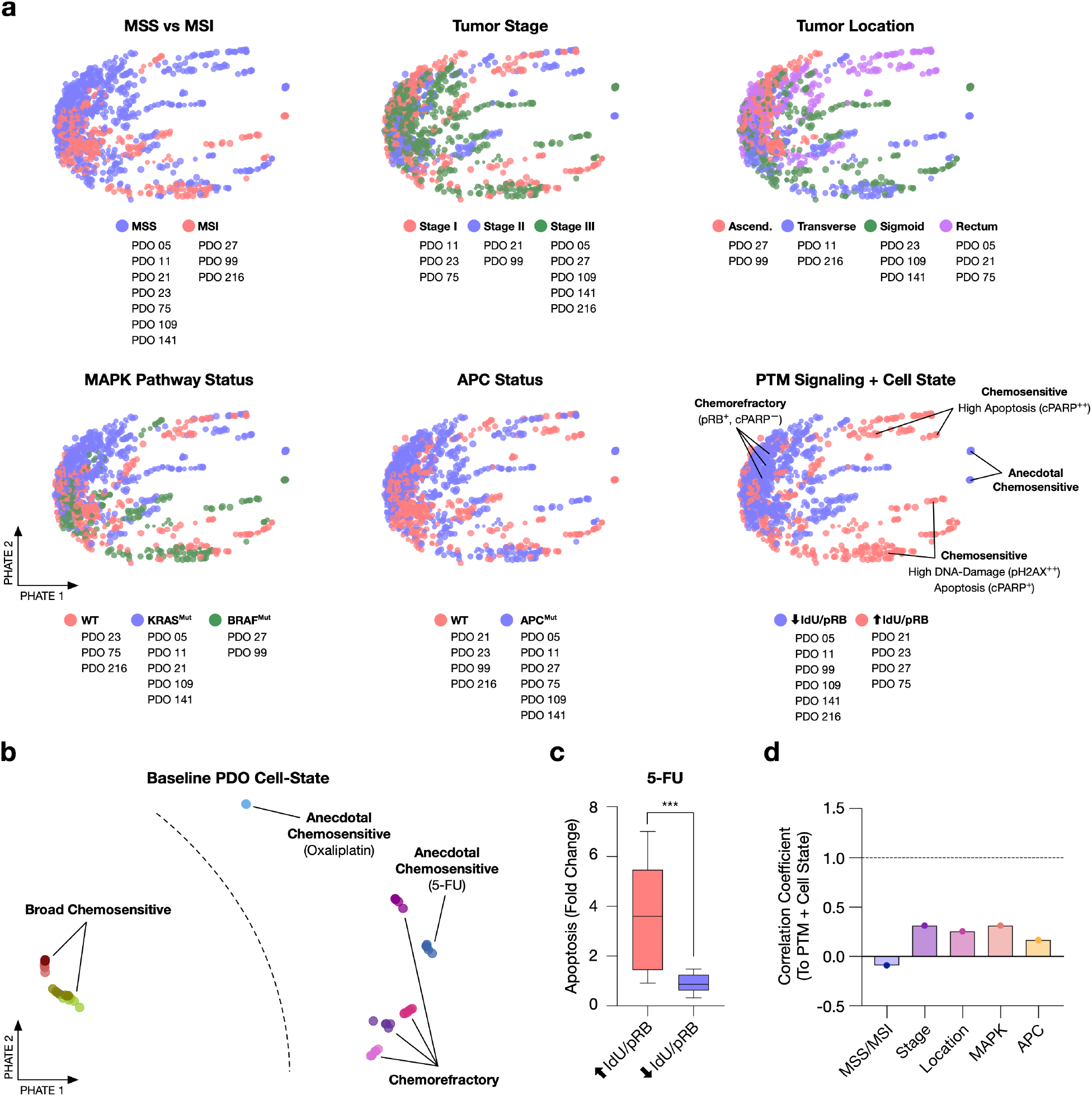
PDO Metadata. **a)** Trellis-PHATE plots of patient metadata. Patient-specific treatment effects do not align with MSS/MSI, tumor stage, tumor location, MAPK pathway mutations, or APC mutations. High baseline cell-cycle activity correlates with broad chemosensitivity. **b)** Trellis-PHATE of baseline PDOs annotated by chemosensitivity. **c)** Quantification of 5-FU chemocytotoxicity in low and high cycling PDOs. **d)** Quantification of the correlation between PDO metadata information and PDO cell-state. Unpaired t-test, *** <0.0001.

**Figure S10:**
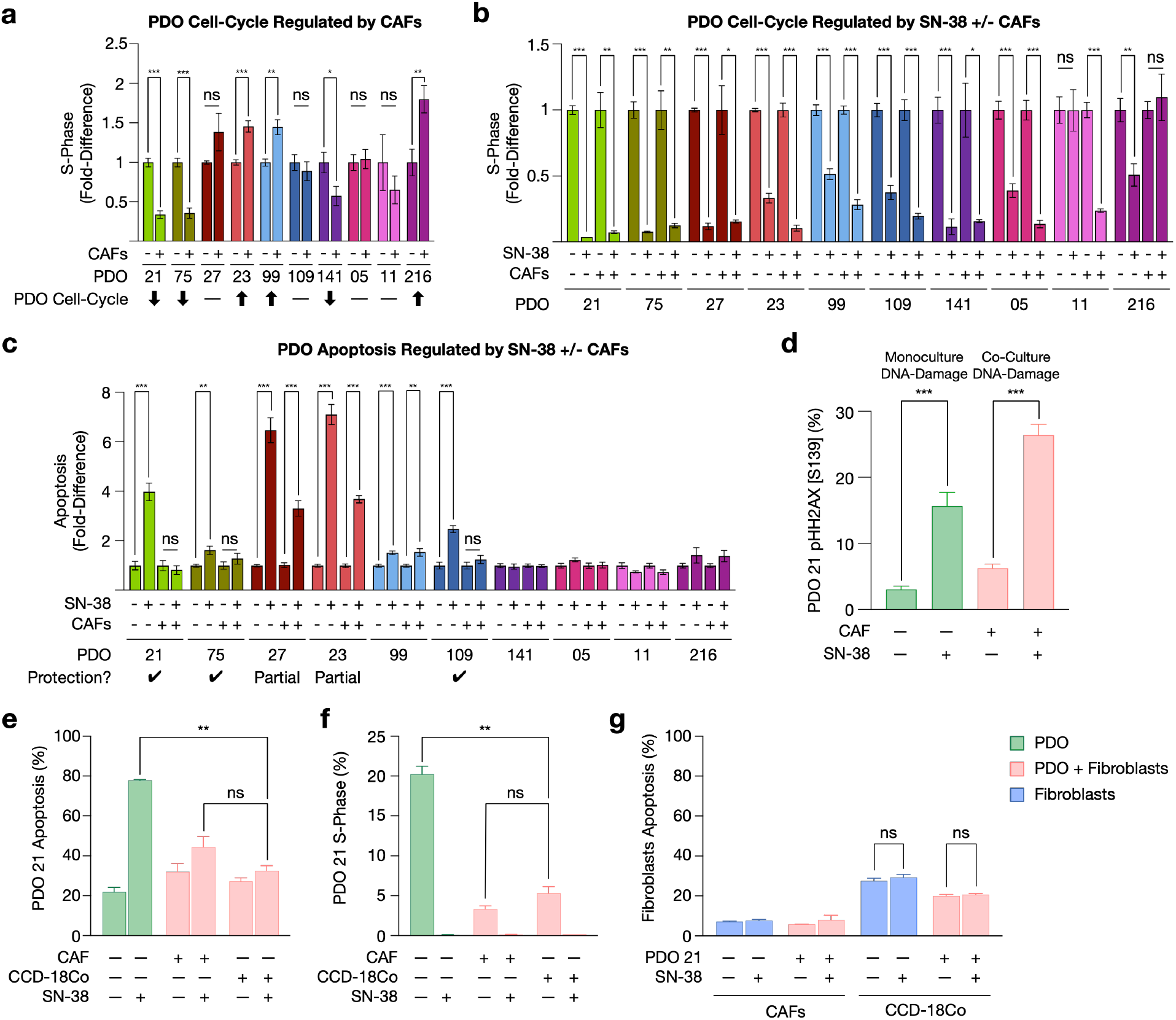
CAF-Induced PDO Cell-State Shifts. **a)** Fold-difference to monoculture of PDO cells in S-phase when co-cultured with CAFs. CAFs both decrease and increase PDO S-phase in a patient-specific manner. **b)** Fold-difference to vehicle controls of PDO cells in S-phase when treated with SN-38 either as PDOs alone or in co-culture with CAFs. CAFs do not alter SN-38 on-target S-phase blockage. **c)** PDO SN-38 induced apoptosis +/- CAFs. Partial CAF-protection is defined as a reduction drug-induced apoptosis in co-culture relative to monoculture, yet still apoptosis is still > 1.5-fold over control and statistically significant. **d)** SN-38 induces on-target DNA-double strand breaks (DSB) (pHH2AX^+^) in PDO 21 irrespective of CAFs. **e-g)** PDO 21 chemoprotection via CCD-18Co normal colon fibroblasts. Unpaired t-test, *** <0.0001, ** <0.001, * <0.01. ns, not significant.

**Figure S11:**
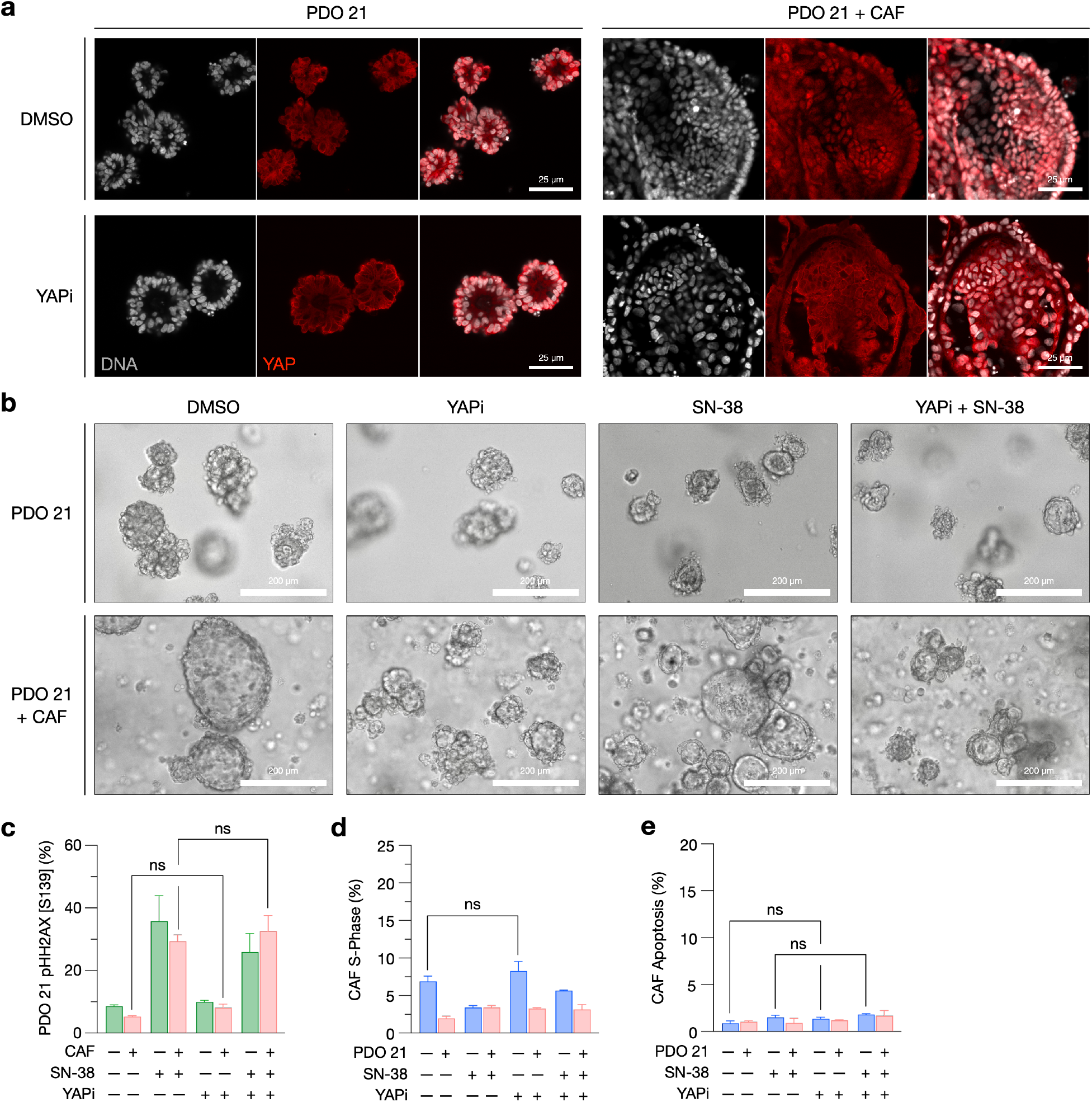
YAP Inhibition of CAF-induce Chemoprotection. **a)** CAF-induced nuclear translocation of YAP (red) to PDO nucleus (white) is reversed by Verteporfin (YAPi). Scale bar = 25 *μ*m. **b)** PDO 21 morphology +/- CAFs, +/- YAPi, +/- SN-38. YAPi reverses CAF-induced cyst-like morphology. Scale bar = 200 *μ*m. **c)** YAPi does not alter SN-38 induces on-target DNA-double strand breaks (pHH2AX^+^) in PDOs. **d)** YAPi does not alter S-phase or **e)** apoptosis in CAFs. Unpaired t-test, *** <0.0001, ** <0.001, * <0.01. ns, not significant.

